# NF-κB Signaling is Required for X-Chromosome Inactivation Maintenance Following T cell Activation

**DOI:** 10.1101/2024.02.08.579505

**Authors:** Katherine S. Forsyth, Natalie E. Toothacre, Nikhil Jiwrajka, Amanda M. Driscoll, Lindsey A. Shallberg, Charlotte Cunningham-Rundles, Sara Barmettler, Joceyln Farmer, James Verbsky, John Routes, Daniel P. Beiting, Neil Romberg, Michael J. May, Montserrat C. Anguera

## Abstract

X Chromosome Inactivation (XCI) is a female-specific process which balances X-linked gene dosage between sexes. Unstimulated T cells lack cytological enrichment of *Xist* RNA and heterochromatic modifications on the inactive X chromosome (Xi), and these modifications become enriched at the Xi after cell stimulation. Here, we examined allele-specific gene expression and the epigenomic profiles of the Xi following T cell stimulation. We found that the Xi in unstimulated T cells is largely dosage compensated and is enriched with the repressive H3K27me3 modification, but not the H2AK119-ubiquitin (Ub) mark, even at promoters of XCI escape genes. Upon CD3/CD28-mediated T cell stimulation, the Xi accumulates H2AK119-Ub and H3K27me3 across the Xi. Next, we examined the T cell signaling pathways responsible for Xist RNA localization to the Xi and found that T cell receptor (TCR) engagement, specifically NF-κB signaling downstream of TCR, is required. Disruption of NF-κB signaling, using inhibitors or genetic deletions, in mice and patients with immunodeficiencies prevents Xist/XIST RNA accumulation at the Xi and alters expression of some X-linked genes. Our findings reveal a novel connection between NF-κB signaling pathways which impact XCI maintenance in female T cells.

## INTRODUCTION

T cells are important adaptive immune cells with many diverse roles including cytolysis of infected cells, the development of a B cell-mediated neutralizing antibody response, and inflammatory cytokine secretion. Immune responses are known to exhibit sex differences^1^ which can impact outcomes following immune challenges^2–5^. While females can exhibit sex-biased protection from a variety of pathogens^1^, they account for the majority of cases for many prevalent autoimmune diseases, including systemic lupus erythematosus (SLE), systemic sclerosis, and multiple sclerosis^6^, where T cells are often dysregulated ^7–9^. The mechanisms underlying sex differences with T cell function and activation, and how these pathways become altered in female-biased autoimmune diseases are not well understood.

T cell activation requires T cell receptor (TCR) engagement of the Major Histocompatibility Complex (MHC) molecule presenting a cognate peptide (signal 1), in combination with co-stimulation through CD28 (signal 2)^10^. Cytokines such as IL-2 present during TCR engagement with MHC can polarize T cell differentiation and subsequent effector functions (signal 3)^10^. There are various signaling cascades which become activated in response to these three critical signals, including canonical NFκB and calcium pathways from signal 1, PI3K/AKT pathways from signal 2, and Jak/Stat pathways from signal 3. Importantly, T cell activation signals can be mimicked *in vitro* with αCD3 and αCD28 antibodies and addition of specific cytokines.

The X chromosome is enriched for genes with immune functions, and the expression of these genes is carefully regulated to ensure cell viability and function^6,11^. T cell function is influenced by various X chromosome-encoded genes, including *FOXP3, CD40L*, *NEMO*, and *CXCR3*. Female eutherian mammals regulate X-linked gene expression through the process of X Chromosome Inactivation (XCI), where either the paternally- or maternally-inherited X chromosome is randomly chosen for silencing in each cell. XCI is initiated early in female embryonic development when the future inactive X Chromosome (Xi) upregulates the long noncoding RNA *Xist*^12,13^. *Xist* RNA subsequently coats the length of the future Xi *in cis*^14–16^ and recruits polycomb repressive complexes (PRC1, PRC2) which add repressive histone modifications H2AK119-ubiquitin (Ub) and H3K27me3^17–19^ across the chromosome. After the formation of the Xi, XCI maintenance is maintained throughout cell division and for the lifetime of the cell, with *Xist* RNA and heterochromatic histone modifications stably associated with the Xi. While the Xi is mostly transcriptionally silent, some genes are expressed by the Xi and the Xa, thereby escaping XCI^20–24^. Some genes that escape XCI have Y-linked homologs (X-Y gene pairs), and XCI escape gene expression can vary by cell type ^20–24^. For example, the X-linked gene *Cxcr3* escapes XCI in stimulated T cells following *Leishmania* infection and correlates with more potent T cell effector function ^25^.

Unlike most somatic cells, T cells from both mice and humans maintain XCI in a dynamic manner whereby *Xist* RNA and repressive histone modifications are cytologically diffuse and not enriched on the Xi in unstimulated T cells, yet *Xist* is robustly transcribed^26,27^. Following stimulation *in vitro* with αCD3/αCD28, *Xist* RNA and heterochromatic histone modifications such as H3K27me3 and H2AK119Ub become enriched at the Xi between 36 and 48 hours of stimulation^26,27^. However, the specific T cell activation signals and downstream signal transduction elements necessary for *Xist* RNA re-localization to the Xi are unknown. Furthermore, whether the Xi remains dosage compensated in unstimulated T cells which lack cytological enrichment of *Xist* RNA and heterochromatic modifications, is not known.

In this manuscript, we utilized allele-specific RNAseq and CUT&RUN approaches to determine the transcriptional and epigenomic profile of the Xi in unstimulated and *in vitro*-activated T cells. We find that the Xi in unstimulated T cells is dosage compensated, with an enrichment of H3K27me3 modifications at specific genes and regions, and that H2K119-Ub progressively accumulates across the Xi after stimulation. We also investigated the signaling requirements necessary for localization of *Xist* RNA and heterochromatic marks to the Xi. Using chemical inhibition, genetic mouse models, and human peripheral blood samples, we discovered that canonical NF-κB signaling is required for *Xist* RNA localization in activated T cells and the regulation of some X-linked genes. Our findings link NF-κB signaling pathways with dynamic XCI maintenance in female T cells.

## RESULTS

### The Xi is dosage compensated in unstimulated T cells lacking cytological epigenetic marks

Unstimulated T cells lack a canonical *Xist* RNA ‘cloud’ and some heterochromatic marks at the Xi^26,27^, suggesting that the Xi might be partially or fully reactivated. To determine whether the Xi in unstimulated T cells is dosage compensated, we quantified allele-specific transcription from each X chromosome using a F1 hybrid mouse model of skewed XCI generated by mating female C57BL/6 mice harboring a heterozygous *Xist* deletion to male *M. m. castaneous* mice (‘F1 mus x cast’). The resulting F1 females harboring the inherited *Xist* deletion therefore exhibit selective XCI of the paternally-derived X chromosome from Cast mice, resulting in skewed XCI. Single nucleotide polymorphisms (SNPs) between the C57BL/6 and Cast genomes can therefore be used to assign reads to the Xi or Xa (**Figure 1A**). We isolated splenic unstimulated CD3^+^ T cells from female F1 mus x cast mice and performed allele-specific RNA sequencing. We detected significantly higher levels of normalized reads (reads per million; RPM) mapping to the Xa compared to the Xi (**Figure 1B**), indicating that the Xi is dosage compensated in unstimulated T cells. We also observed that there is some transcription specifically mapping to the Xi, at approximately 20% of the level of the Xa (**Figure 1B**). For autosomes, we observed a similar number of RPM from each allele (**Figure S1A**). Thus, the Xi is largely transcriptionally silent in unstimulated T cells despite the absence of the canonical *Xist* RNA ‘cloud’ and immunofluorescence enrichment of heterochromatic modifications.

**Figure 1.**
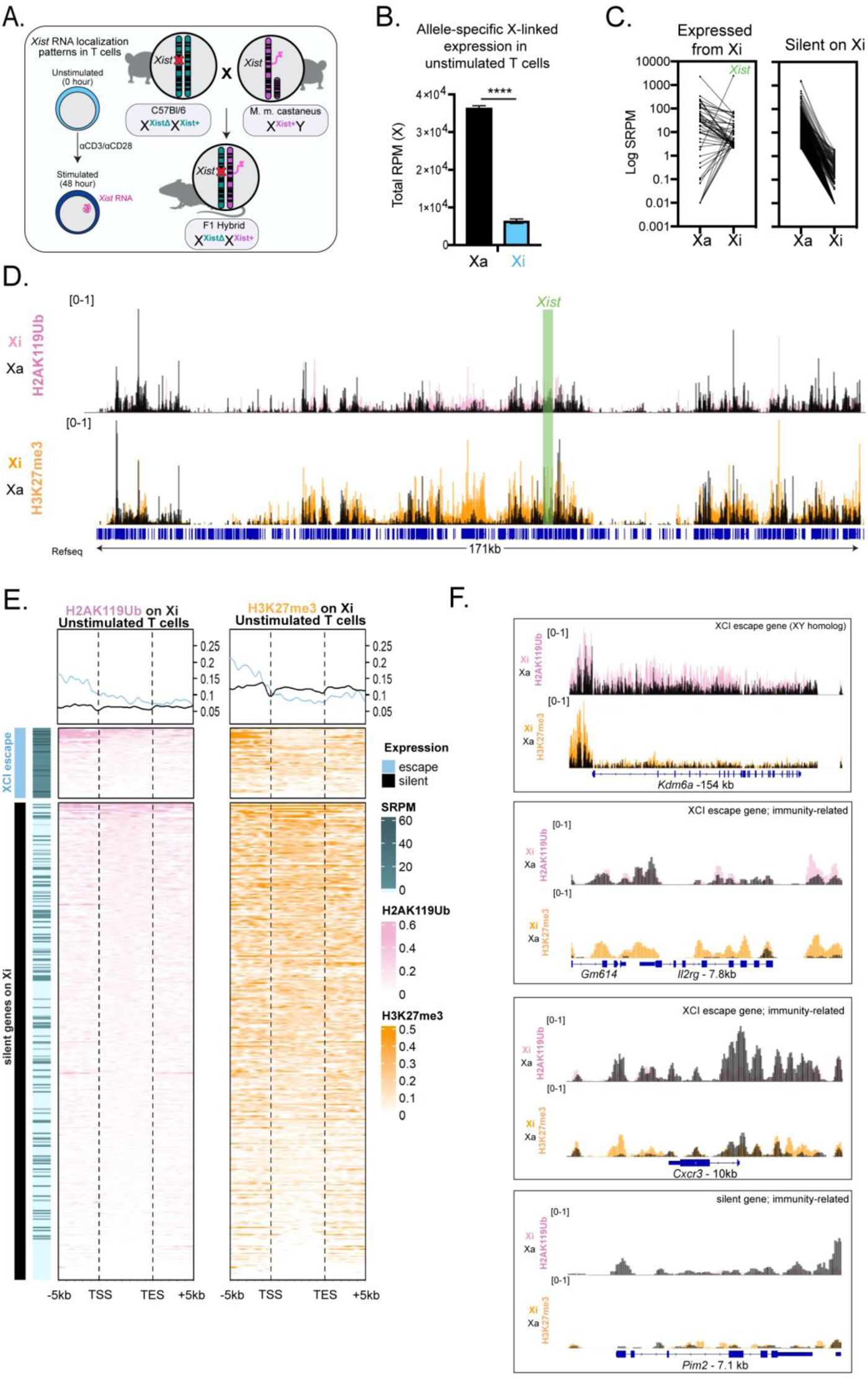
The Xi in unstimulated T cells is dosage compensated with transcription from the Xi and unique epigenetic features at both expressed and silent genes. **A**. Schematic for *Xist* RNA localization patterns in unstimulated and stimulated T cells (left), and breeding setup to generate F1 mus x cast mice for allele-specific RNA-seq and CUT&RUN (right). **B**. Total reads per million (RPM) coming from the Xa and Xi alleles. Averages of 3 biological replicates with SD are shown. Statistical significance quantified using unpaired T test with Welch’s correction, **** p< 0.0001. **C**. Log SRPM from Xa and Xi alleles in unstimulated T cells, with each paired line representing one gene, separated by genes escaping XCI (“expressed from Xi”) and genes subject to Xi (“silent on Xi”). Statistical significance quantified using the Mann-Whitney U test comparing the Xi/(Xa+Xi) SRPM for each gene in expressed and silent categories, ****p<0.0001. **D**. CPM-normalized track of X chromosome wide enrichment of H2AK119Ub (top) and H3K27me3 (bottom) in unstimulated T cells. Black represents enrichment on the Xa, while pink represents Xi enrichment of H2AK119Ub and orange represents Xi enrichment of H3K27me3. Green bar indicates region around *Xist* promoter. **E**. Heatmap of enrichment of H2AK119Ub (left, pink) and H3K27me3 (right, orange) for all expressed (RPKM ≥ 1) X-linked genes, split up by XCI escape status. Metaplots show average enrichments 5kb upstream of the transcription start site (TSS) to 5kb downstream of the transcription end site (TES). SRPM from the Xi for each gene is represented by teal horizontal lines. Blue represents genes escaping XCI and black represents genes subject to XCI. **F**. CPM-normalized tracks of H2AK119Ub (top) and H3K27me3 (bottom) enrichment at individual genes. Black represents enrichment on the Xa, while pink represents Xi enrichment of H2AK119Ub and orange represents Xi enrichment of H3K27me3. The genes shown are *Kdm6a* (154kb), *Il2rg* (7.8kb), *Cxcr3* (10kb), *Pim2* (7.1kb).

To determine the X-linked genes that are expressed from the Xi and therefore ‘escape’ XCI silencing, we used a previously published method^21,28^ to identify XCI escape genes in female murine unstimulated T cells. Using this approach, a gene was considered expressed from the Xi if: (1) the diploid (mus + cast reads) Reads Per Kilobase Million (RPKM) was greater than 1; (2) haploid expression (SNP containing Reads Per Million, SRPM) was greater than 2; and (3) the probability of transcription from the Xi, as determined via a previously published binomial model^21^, was greater than 0. We identified a total of 368 X-linked genes that are expressed in unstimulated T cells, and 51 of these were expressed from the Xi (**Supplemental Table S1, Supplemental Table S2**). Unlike *Xist*, which is exclusively expressed from the Xi, some XCI escape genes are more likely to have similar expression levels from the Xi and Xa (**Figure 1C**). We identified some novel T cell specific XCI escape genes with known roles in T cell function, including *Cxcr3, Il2rg, Cul4b,* in addition to the X-Y gene pairs known to escape XCI in other cell types (*Kdm6a, Ddx3x),* and *Ftx* which is an activator of *Xist* expression (**Table S1**). Thus, while unstimulated T cells are mostly dosage compensated, there are 51 genes that are expressed from the Xi.

### The Xi in unstimulated T cells retains enrichment of H3K27me3 modifications across both silent and expressed genes

Cytological analyses of the Xi using immunofluorescence indicates that the Xi in unstimulated T cells lacks enrichment of the heterochromatic modifications H3K27me3 and H2AK119-ubiquitin (Ub)^27,29^. Given our observations that the Xi in unstimulated T cells is largely dosage compensated, we asked whether these repressive histone modifications were indeed absent across the Xi in unstimulated T cells, at higher resolution. We performed allele-specific CUT&RUN for H2AK119-Ub and H3K27me3 in unstimulated splenic CD3^+^ T cells from female F1 mus x cast mice (**Figure 1A**). We observed low levels of H2AK119-Ub across the Xi, at levels comparable to the Xa (**Figure 1D**, top track in pink for Xi, black for Xa). Surprisingly, there were high levels of H3K27me3 enrichment across the Xi, at levels higher compared to the Xa (**Figure 1D**). Examination of 5kb regions flanking transcriptional start sites (TSS) across the Xi confirmed that there are higher levels of H3K27me3 than H2AK119-Ub. Similarly, examination of Xi gene bodies also revealed more H3K27me3 than H2AK119-Ub irrespective of gene expression (**Figure 1E**). For regions upstream of the transcription start site (TSS), X-linked genes expressed from the Xi (XCI ‘escape’; blue line) have more H3K27me3 and H2AK119-Ub compared to silent genes (black line) (**Figure 1E**). Most XCI escape genes have less H3K27me3 at gene bodies compared to silent genes (**Figure 1E**). We identified a subset of XCI escape genes (*Ddx3x, Eif2s3x, Ftx, Kdm5c, Kdm6a, Ogt, Tbl1x, Xkrx*) which have increased levels of H3K27me3 at regions upstream of their promoters. For example, *Kdm6a* and *Eif2sx* have significant Xi-specific enrichment of H3K27me3 at upstream promoter regions and reduced levels across the gene bodies, yet there is also Xi-specific enrichment of H2AK119-Ub at both regions (**Figure 1F and Supplemental Figure 1B**). The immunity-related X-linked gene *Cxcr3* (expressed from Xi) has more Xi-specific H3K27me3 marks upstream and downstream of the gene body compared to the Xa, yet H2AK119-Ub levels are similar for Xa and Xi (**Figure 1F**). Another immunity-related X-linked gene, *Il2rg (*expressed from Xi), also has high levels of H3K27me3 upstream and across the gene body on the Xi compared to the Xa, and high levels of H2AK119-Ub at the Xi upstream promoter region (**Figure 1F**). *Pim2*, a serine/threonine kinase that functions in signal transduction for T cell growth and proliferation, is repressed on the Xi in unstimulated cells, and has low levels of H3K27me3 and H2AK119-Ub on the Xi, similar to the Xa (**Figure 1F**). *Huwe1*, which is transcriptionally silent from the Xi in unstimulated T cells (**Table S2**) has elevated levels of H3K27me3 across the gene body (bottom track), and low levels of H2AK119-Ub on the Xi, similar to the Xa (**Supplemental Figure 1B**). Taken together, these findings reveal that there are diverse patterns of H3K27me3 enrichment for both XCI escape and silent genes on the Xi in unstimulated T cells.

### XCI escape genes in stimulated T cells with a dosage compensated Xi

Following TCR engagement, T cells exit quiescence and re-enter the cell cycle with the first cell division occurring at 48 hrs^30^. We also performed allele-specific RNA sequencing on splenic CD3^+^ T cells from F1 mice stimulated with αCD3/αCD28 for 48 hours *in vitro*. We observed more transcription from the Xa compared to the Xi indicative of dosage compensation (**Figure 2A**) similar to unstimulated T cells, and autosomes have similar RPM values for paternal and maternal alleles (**Supplementary Figure S2A**). While the Xi is mostly transcriptionally silent in stimulated T cells, there is transcription from the Xi (blue bar, **Figure 2A**), at similar levels compared to unstimulated T cells (**Figure 1B**). Using our Escape_Gene_Calculator, we identified 332 genes expressed from the X in stimulated T cells, and 35 of these are significantly expressed from the Xi (**Supplemental Table S1, Supplemental Table S2**). Most XCI escape genes expressed in stimulated T cells were expressed at roughly similar levels between the Xa as the Xi (except for *Xist* and 7 other genes) (**Figure 2B**). Comparing the XCI escape genes between unstimulated (51 X-linked genes) and stimulated T cells (35 X-linked genes), the majority of XCI escape genes (31 X-linked genes) are expressed from the Xi in T cells regardless of stimulation state (**Figure 2C and Supplemental Table S1**). We analyzed stimulation-dependent changes in XCI escape genes and identified 20 XCI escape genes specific to unstimulated T cells, and 4 escape genes specific to stimulated T cells: *Gk, Gm14539, Gm15190*, and *Pim2* (**Figure 2C**). Most X-Y pair genes escape XCI regardless of stimulation, and there were two genes that escaped XCI specifically in unstimulated T cells: *Tbl1x* and *Xkrx* (**Table S1**). Some X-linked genes are expressed from the Xi in both unstimulated and stimulated T cells (*Gm6472, Il2rg, Med14, Cul4b* and *Kdm6a*), while others are repressed in stimulated T cells (*Cxcr3* and *Mtmr1)* (**Supplementary Table S1**). Thus, T cells exhibit cell-specific XCI escape genes that are mostly similar across stimulation state.

**Figure 2.**
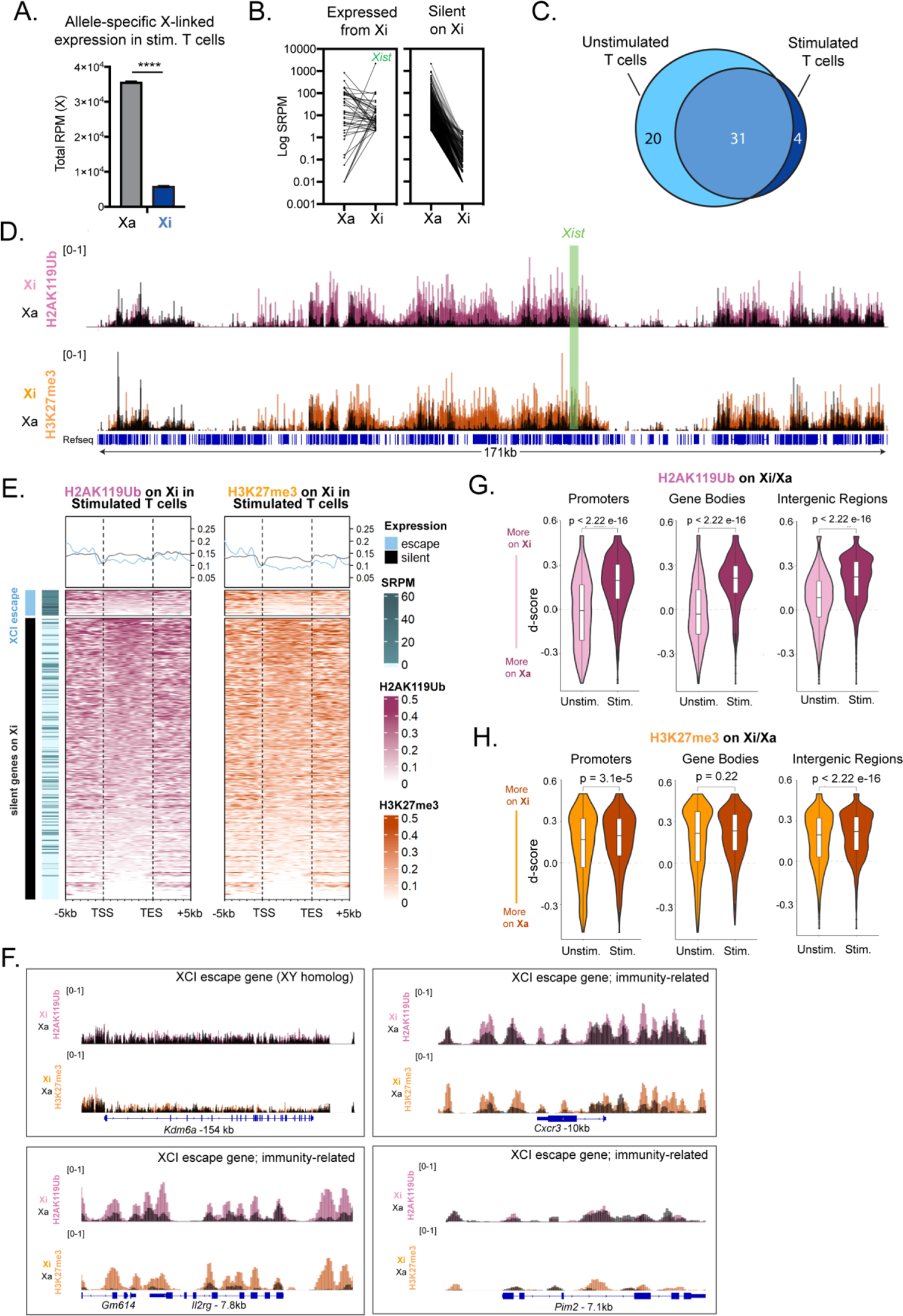
The Xi in stimulated T cells acquires H2AK119-Ub marks and XCI escape genes have distinct epigenetic features. **A.** Total reads per million (RPM) coming from the Xa and Xi alleles. Averages of 3 biological replicates with SD are shown. Statistical significance quantified using unpaired T test with Welch’s correction, **** p< 0.0001 **B.** Log SRPM from Xa and Xi alleles in stimulated T cells, with each paired line representing one gene, separated by genes escaping XCI (“expressed from Xi”) and genes subject to Xi (“silent on Xi”). Statistical significance quantified using the Mann-Whitney U test comparing the Xi/(Xa+Xi) SRPM for each gene in expressed and silent categories, ****p<0.0001. **C.** Proportional Venn diagram showing overlap of XCI escape genes between unstimulated T cells (light blue) and stimulated T cells (dark blue). **D.** CPM-normalized track of X chromosome wide enrichment of H2AK119Ub (top) and H3K27me3 (bottom) in unstimulated T cells. Black represents enrichment on the Xa, while dark pink represents Xi enrichment of H2AK119Ub and dark orange represents Xi enrichment of H3K27me3. Green bar indicates region around *Xist* promoter. **E.** Heatmap of enrichment of H2AK119Ub (left, dark pink) and H3K27me3 (right, dark orange) for all expressed (RPKM ≥ 1) X-linked genes, split up by XCI escape status. Metaplots show average enrichments 5kb upstream of the transcription start site (TSS) to 5kb downstream of the transcription end site (TES). SRPM from the Xi for each gene is represented by teal horizontal lines. Blue represents genes escaping XCI and black represents genes subject to XCI. **F.** CPM-normalized tracks of H2AK119Ub (top) and H3K27me3 (bottom) enrichment at individual genes. Black represents enrichment on the Xa, while dark pink represents Xi enrichment of H2AK119Ub and dark orange represents Xi enrichment of H3K27me3. The genes shown are *Kdm6a* (154kb), *Il2rg* (7.8kb), *Cxcr3* (10kb), *Pim2* (7.1kb). **G.** D-score analysis of H2AK119Ub accumulation at promoter, gene body, and intergenic regions. Unstimulated T cells are in pink and stimulated T cells are in dark pink. Biological replicates (n=3) were averaged for each timepoint. P-values were calculated using a Wilcoxon signed-rank test with Benjamini Hochberg correction. **H.** D-score analysis of H3K27me3 accumulation at promoter, gene body, and intergenic regions. Unstimulated T cells are in orange and stimulated T cells are in dark orange. Biological replicates (n=3) were averaged for each timepoint. P-values were calculated using a Wilcoxon signed-rank test with Benjamini Hochberg correction.

### T cell stimulation increases H2AK119-Ub across the Xi, resulting in gene-specific epigenomic profiles for XCI escape genes

Using alle-specific profiling, we investigated how T cell stimulation impacts enrichment of heterochromatic histone tail modifications across the Xi. We performed CUT&RUN for H2AK119-Ub and H3K27me3 modifications using *in vitro* stimulated CD3^+^ T cells from female F1 mus x cast mice. The Xi in stimulated T cells is enriched for both H2AK119-Ub and H3K27me3 modifications, at levels higher than the Xa (**Figure 2D**). Gene bodies and 5kb flanking regions across the Xi in stimulated T cells have similar levels of enrichment of H3K27me3 than H2AK119-Ub for silent genes (black lines, **Figure 2E**). Intriguingly, XCI escape genes have lower levels of H3K27me3 across Xi gene bodies in stimulated T cells, yet H2AK119-Ub levels are relatively similar (**Figure 2E**). We observed similar enrichment levels of H3K27me3 and H2AK119-Ub at promoter regions of the Xi for XCI escape and silent X-linked genes in stimulated T cells (**Figure 2E**). Examination of heterochromatic histone modifications for XCI escape genes revealed distinct profiles depending on the class of XCI escape. For XCI escape genes with Y-linked homologs (*Kdm6a, Eif2s3x*), the Xi and Xa in stimulated T cells have similar enrichment of H3K27me3 and H2AK119-Ub modifications (**Figure 2F** and **Supplemental Figure 2B**). However, for immunity-related XCI escape genes expressed in stimulated T cells (*Il2rg, Pim2*), the Xi has higher levels of H3K27me3 and H2AK119-Ub compared to the Xa, across these gene regions (**Figure 2F**), representing specific epigenetic signatures for gene reactivation in a cell type/immune context. For a silent, non-immune X-linked gene (*Huwe1*), the Xi had much higher enrichment of both H3K27me3 and H2AK119-Ub compared to the Xa, surpassing the levels observed for immunity-related genes (**Supplemental Figure 2B**).

Next, we compared the enrichment of each modification on the Xi relative to the Xa, between unstimulated and stimulated T cells. Across the Xi, we observed that the Xi gains higher levels of H2AK119-Ub with stimulation, while levels of H3K27me3 remain constant with stimulation (**Supplemental Figure 2C** and **Supplemental Figure 2D**). To quantify enrichment changes at specific regions across the Xi (promoters, gene bodies, intergenic regions), we calculated the d-score for the ratio of Xi and Xa enrichment for each modification, where positive values reflect higher levels on the Xi and negative d-score values reflect enrichment on the Xa. We found that upon T cell stimulation, there are significant increases in H2AK119-Ub enrichment on the Xi specifically at promoters, gene bodies, and intergenic regions (**Figure 2G**). For H3K27me3, the Xi in unstimulated T cells has elevated levels of this mark at promoters, gene bodies, and intergenic regions, and stimulation increases enrichment specifically at promoters and intergenic regions on the Xi (**Figure 2H**). Promoter regions of XCI escape genes expressed in both unstimulated and stimulated T cells have higher levels of both H2AK119-Ub and H3K27me3 compared to silent genes on the Xi (red lines compared to black lines; **Supplemental Figure 2E**). Silent X-linked genes on the Xi have high levels of H2AK119-Ub which do not change across the +/-5kB gene regions for both unstimulated and stimulated T cells, and have an increase in levels of H2AK119-Ub during stimulation (black lines; **Supplemental Figure 2E**). Together, these data demonstrate that T cell stimulation changes the epigenetic environment across the Xi, with gene-specific enrichment of modifications depending on expression status of the gene.

### T cell receptor signaling is necessary to localize Xist RNA and H2AK119-Ub to the Xi

We have previously demonstrated that T cell stimulation with αCD3/αCD28 results in the re-localization of *Xist* RNA and heterochromatic modifications to the Xi^26,27^, yet the specific T cell signaling requirements for localizing these epigenetic marks are unknown. We asked whether T cell receptor (TCR) engagement is necessary for localization by stimulating cells *in vitro* with either αCD3 alone or cognate peptide-MHC complexes (**Figure 3A**) and performing sequential *Xist* RNA FISH and IF for H2AK119-Ub. Stimulating T cells with αCD3 alone is sufficient for *Xist* RNA ‘clouds’ at levels slightly less than stimulation with αCD3/αCD28, and similar proportions of nuclei with H2AK119-Ub foci are observed for αCD3 alone and αCD3/αCD28 (**Figure 3B-C**). To determine whether TCR signal strength influences *Xist* RNA and H2AK119-Ub localization to the Xi, we isolated splenic T cells from female transgenic mice that express the OT-I TCR specific for the H2-K^b^-restricted SIINFEKL peptide^31^. We peptide-pulsed splenocytes with either the SIINFEKL peptide or peptides that have single amino acid substitutions with decreased affinity for the OT-I TCR^32^ (**Figure 3A**), then co-cultured the peptide-pulsed splenocytes with T cells from congenic OT-I TCR transgenic mice for 72hrs and sorted the OT-I T cells (**Figure S3**) for sequential *Xist* RNA FISH and H2AK119Ub. We observed robust *Xist* RNA ‘clouds’ and co-localization of H2AK119-Ub foci in OT-I T cells co-cultured with the cognate SIINFEKL peptide (**Figure 3D-E**). T cells cultured with the two peptide derivatives with intermediate TCR affinity (SII<u>V</u>FEKL and SII<u>Q</u>FEKL) have similar levels of *Xist* RNA localization and H2AK119-Ub foci as SIINFEKL co-cultures (**Figure 3D-E**). However, T cells cultured with the peptide derivative with lowest TCR affinity (<u>E</u>IINFEKL) have minimal *Xist* RNA ‘clouds’ and H2AK119-Ub foci (**Fig. 3D-E**). These data demonstrate that TCR stimulation is necessary for localizing Xist RNA and heterochromatin marks to the Xi, and suggest that there is a minimum threshold of TCR affinity of peptide-MHC required for *Xist* RNA and heterochromatic mark localization, independent of TCR signaling strength.

**Figure 3.**
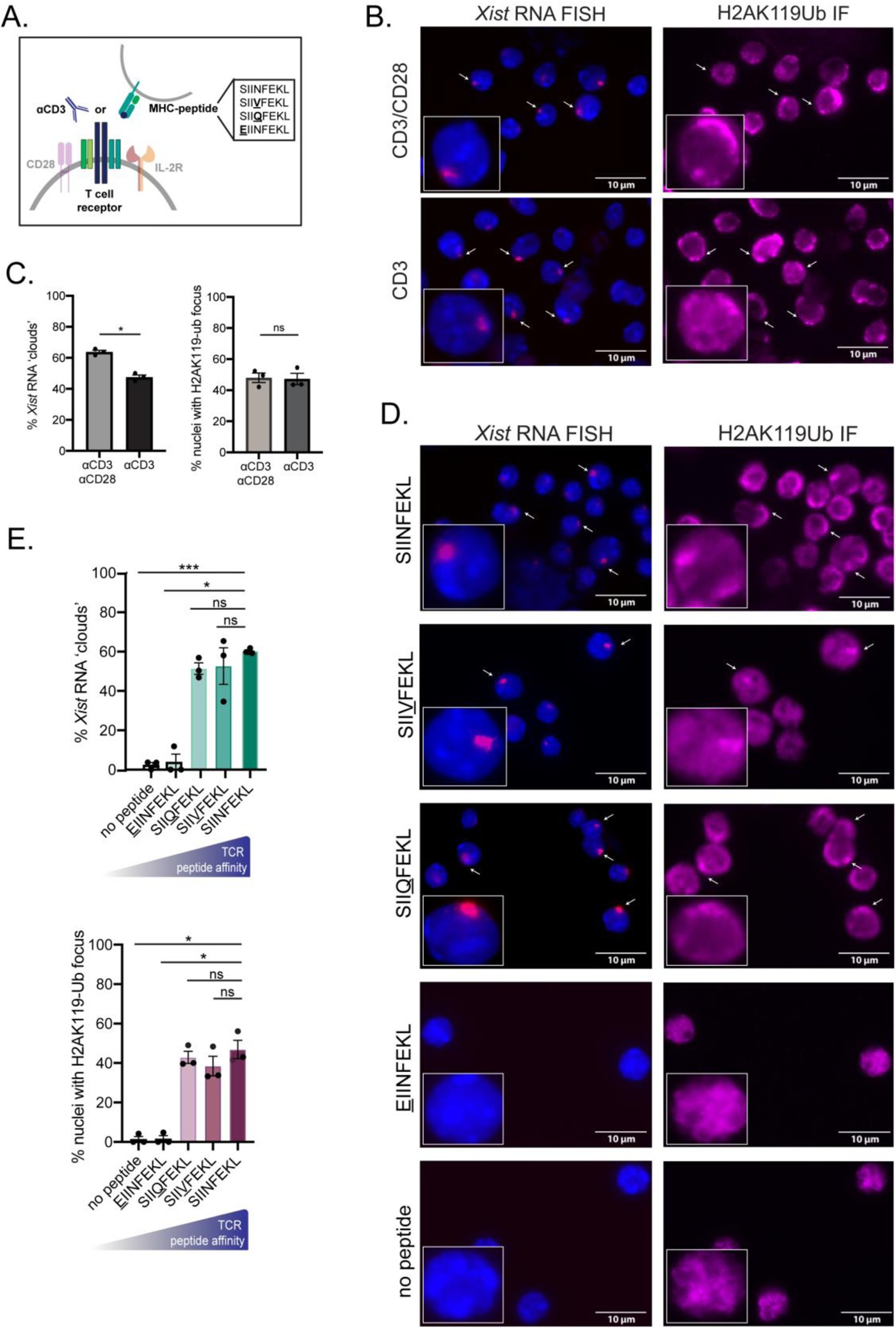
TCR signaling, independent of signal strength, is necessary for Xist RNA and H2AK119-Ub localization to the Xi. **A**. Schematic of T cell activation involving the T cell receptor, mediated by either an αCD3 antibody or MHC-peptide complexes. The amino acid sequences of the peptides with various affinities for the OT-I T cell receptor are shown. **B**. Representative fields (from 1 experiment) showing sequential *Xist* RNA FISH and H2AK119-Ub immunofluorescence (IF) for T cells activated with both αCD3/αCD28 (top) or αCD3 alone (bottom) for 72 hours. **C**. Quantification of *Xist* RNA clouds (left) and H2AK119-Ub foci for T cells cultured with either αCD3/αCD28 or αCD3 alone for 72 hours. Averages of 3 independent experiments with SEM are shown. Statistical significance quantified using unpaired T test with Welch’s correction, * p< 0.05. 102-157 nuclei (FISH) and 85-111 nuclei (IF) were counted per condition per experiment. **D**. Representative fields (from 1 experiment) showing sequential *Xist* RNA FISH and H2AK119-Ub immunofluorescence (IF) for OT-I transgenic T cells cultured with congenic splenocytes peptide-pulsed with either SIINFEKL, SIIVFEKL, SIIQFEKL, EIINFEKL or no peptide for 72 days prior to sorting. **E**. Quantification of *Xist* RNA clouds (top) and H2AK119-Ub foci (bottom) for OT-I transgenic T cells cultured with congenic splenocytes peptide-pulsed with either SIINFEKL, SIIVFEKL, SIIQFEKL, EIINFEKL for 72 hours prior to sorting. Averages of 3 independent experiments with SEM are shown. Statistical significance quantified using one way ANOVA followed by multiple comparisons to anti-SIINFEKL condition, * p<0.05 ** p<0.01, *** p<0.001. 27-144 nuclei (FISH) and 20-106 nuclei (IF) were counted per condition per experiment.

### CD28, IL-2, and calcium signaling and progression through the cell cycle are independent of Xist RNA and H2AK119-Ub localization to the Xi

Co-stimulation through CD28 engagement and cytokine signaling are also important for T cell activation^33,34^. We investigated whether CD28 and IL-2 signaling are necessary for localizing *Xist* RNA and H2AK119-Ub to the Xi (**Figure 4A**). We cultured splenic T cells from wildtype female mice with either αCD28 or soluble IL-2 for 72 hours, then performed sequential *Xist* RNA FISH and IF. Neither αCD28 or soluble IL-2 alone was sufficient to localize *Xist* RNA transcripts to the Xi, and very few H2AK119-Ub foci were detected (**Figure 4B-C**). Next, we asked whether *Xist* RNA and H2AK119-Ub localization correlates with a particular phase of the cell cycle. We cultured splenic T cells with αCD3/αCD28 for 72 hours with various inhibitors known to block cell cycle progression at G0/G1 (rapamycin), S-phase (hydroxyurea), and G2/M phase (nocodozole)^35^, and then isolated cells for sequential *Xist* RNA FISH and IF. We observed similar levels of *Xist* RNA localization and H2AK119-Ub foci across all phases of the cell cycle in arrested T cells as compared to untreated stimulated T cells (**Figure 4D-E**). Thus, T cell cycle progression does not correlate with the changes in *Xist* RNA and H2AK119-Ub localization to the Xi.

**Figure 4.**
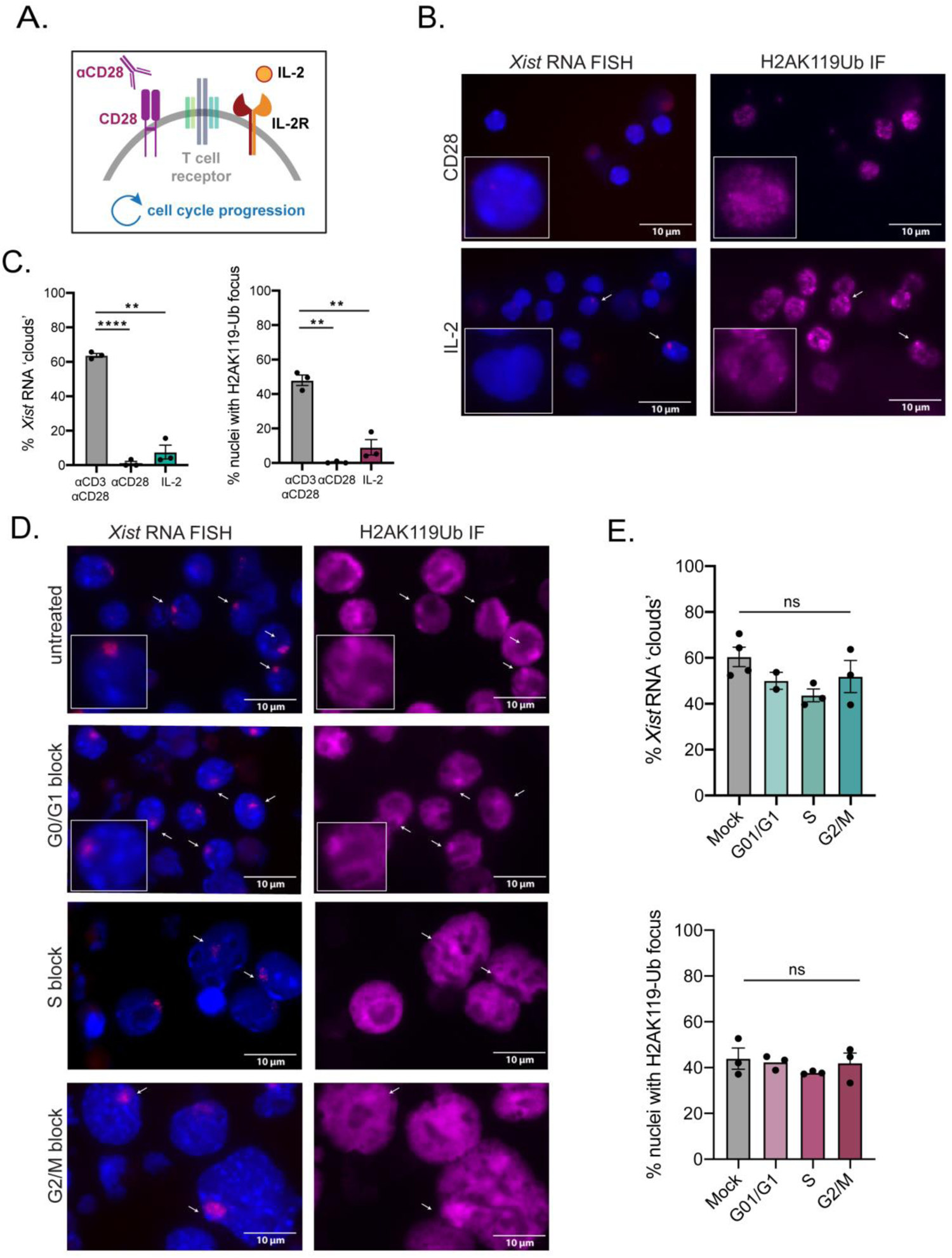
Signaling through CD28 or IL-2 alone are not sufficient for *Xist* RNA and H2AK119-Ub localization to the Xi. **A**. Schematic of CD28 receptor at the T cell surface which is activated by αCD28 antibody; the IL-2 receptor is activated by soluble recombinant IL-2. **B.** Representative fields (from 1 experiment) showing sequential *Xist* RNA FISH and H2AK119-Ub IF for T cells activated with either αCD28 alone (top) or IL-2 alone (bottom) for 72 hours. **C**. Quantification of *Xist* RNA clouds (left) and H2AK119-Ub foci for T cells cultured with either αCD3/αCD28, αCD28 alone, or IL-2 alone for 72 hours. Averages of 3 independent experiments with SEM are shown with each dot representing a biological replicate. Statistical significance quantified using one way ANOVA followed by multiple comparisons to αCD3/αCD28 condition, ** p<0.01, **** p<0.0001. 74-157 nuclei (FISH) and 58-111 nuclei (IF) were counted per condition per experiment. **D.** Representative fields (from 1 experiment) showing sequential *Xist* RNA FISH and H2AK119-Ub IF of αCD3/αCD28 stimulated T cells treated with 100nM rapamycin (G0/G1 block), 200μM hydroxyurea (S block), 1μg/mL nocodozole (G2/M block), or a mock treatment control for 72 hours. **E**. Quantification of *Xist* RNA clouds (top) and H2AK119-Ub foci for αCD3/αCD28 stimulated T cells treated with 100nM rapamycin (G0/G1 block), 200μM hydroxyurea (S block) 1μg/mL nocodozole (G2/M block), or a mock treatment control for 72 hours. Averages of 3 independent experiments with SEM are shown with each dot representing a biological replicate. Statistical significance quantified using one way ANOVA followed by multiple comparisons to mock treatment. 85-135 nuclei (FISH) and 52-146 nuclei (IF) were counted per condition per experiment.

### Canonical NFκB signaling is required for localizing Xist RNA and H2AK119-Ub to the Xi

Because TCR engagement is required for *Xist* RNA and H2K119-Ub modifications to accumulate at the Xi, we investigated downstream pathways of the TCR. After TCR engagement, T cells internalize extracellular calcium through transmembrane channels, as calcium functions as an intracellular signaling molecule for the activation of target enzymes and transcription factors including nuclear factor of activated T cells (NFAT)^36^.

We cultured splenic T cells with αCD3/αCD28 in the absence of exogenous calcium for 72 hours and performed sequential *Xist* RNA FISH and IF for H2AK119-Ub. We found that the absence of extracellular calcium did not significantly change the levels of *Xist* RNA clouds or percentages of H2AK119-Ub foci at the Xi compared to calcium-replete cultured T cells (**Supplemental Figure S4A, B**).

Next, we asked whether the canonical NF-κB pathway, which is downstream of TCR engagement, is involved in re-localizing *Xist* RNA and H2AK119Ub foci to the Xi in T cells. After TCR engagement, the IKK complex consisting of the kinases IKKα and IKKβ, and the scaffold protein NEMO, is activated, and this in turn phosphorylates members of the inhibitory IκB family of proteins resulting in their degradation by the proteasome. IκB degradation releases canonical REL homology domain (RHD)-containing NF-κB protein heterodimers that translocate to the nucleus to drive gene activation^37,38^ (**Figure 5A**). To block canonical NF-κB-mediated gene activation, we used two distinct chemical inhibitors of the IKKβ kinase to prevent IκB phosphorylation: IMD-0354 and TPCA-1^39–41^ (**Figure 5A**). We stimulated T cells with αCD3/αCD28 for 48hrs in the presence of either IMD-0354 or TPCA-1, then collected cells for *Xist* RNA FISH and IF for H2AK119-Ub and H3K27me3, another heterochromatic modification enriched on the Xi in stimulated T cells ^26,27^ (**Figure 2D**). Surprisingly, we observed significant reductions with *Xist* RNA transcripts clustered at the Xi when T cells are stimulated in the presence of either IMD-0354 or TPCA-1, and the effect was concentration dependent (**Figure 5B-C**). The loss of *Xist* RNA localization was independent of both H2AK119-Ub and H3K27me3 enrichment at the Xi, as both modifications were detected as focal enrichments in nuclei lacking canonical *Xist* RNA ‘clouds’ (**Figure 5D-E**). IMD-0354 treatment did not change the number of X chromosomes in stimulated T cells, as 2 Xs were observed in treated cells lacking *Xist* RNA ‘clouds’ and H2AK119-Ub foci colocalized with one X chromosome in sequential RNA FISH/IF/DNA FISH experiments (**Supplemental Figure 4C**). Treatment of immortalized female mouse embryonic fibroblasts (MEFs) with IMD-0354 did not affect *Xist* RNA localization (**Supplemental Figure 4D**), suggesting that NF-κB-mediated impact on *Xist* RNA at the Xi is lymphocyte-specific.

**Figure 5.**
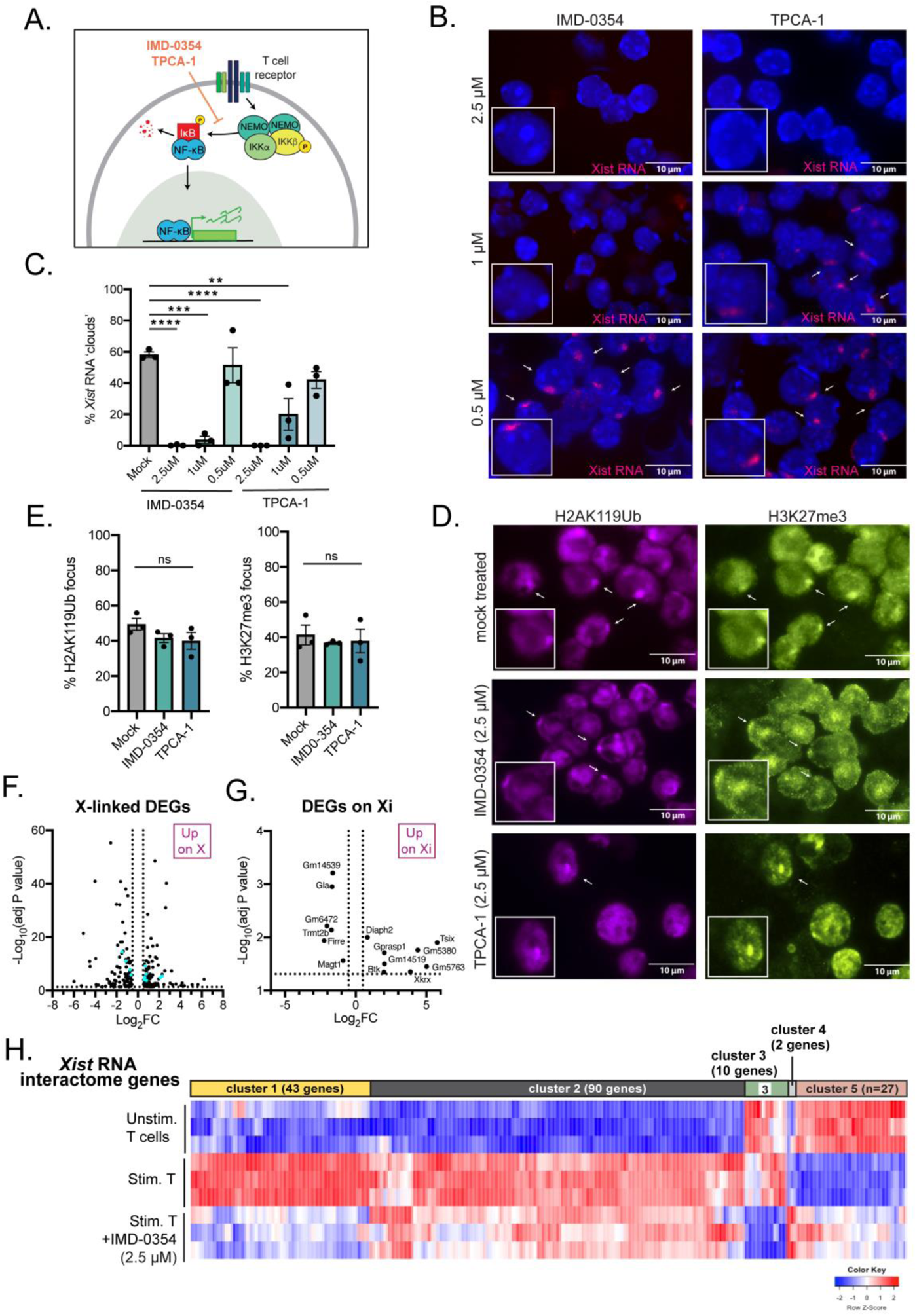
Canonical NF-κB signaling stimulates the re-localization of *Xist* RNA and heterochromatic histone modifications to the Xi through gene activation of *Xist* RNA ‘Interactome’ genes. **A.** Schematic of the canonical pathway of NF-κB activation, where TCR engagement induces phosphorylation of IκB (red) by the IKK complex composed of IKKα, IKKβ, and NEMO. Phosphorylation of IκB induces its ubiquitinylation and proteasomal degradation, releasing NFκB dimers to translocate into the nucleus. Kinase inhibitors IMD0354 and TPCA-1 block phosphorylation of IκB. **B**. Representative fields (from 1 experiment) of *Xist* RNA FISH of αCD3/αCD28 stimulated T cells treated with various concentrations (2.5μM, 1μM, 0.5μM) of IKKβ kinase inhibitors (IMD-0354 or TPCA-1) for 48 hours. **C**. Quantification of *Xist* RNA ‘clouds’ of αCD3/αCD28 stimulated T cells treated with various concentrations (2.5μM, 1μM, 0.5μM) of IKKβ kinase inhibitors (IMD-0354 or TPCA-1) for 48 hours. Averages of 3 independent experiments with SEM are shown with each dot representing a biological replicate. Statistical significance quantified using one way ANOVA followed by multiple comparisons to mock treated condition, ** p<0.01, *** p<0.001, **** p<0.0001 with 102-173 nuclei counted per condition per experiment. **D**. Representative fields (from 1 experiment) of H2AK119-Ub IF (left) and H3K27me3 IF (right) of αCD3/αCD28 stimulated T cells treated with 2.5μM of IKKβ kinase inhibitors IMD-0354 or TPCA-1 for 48 hours. **E.** Quantification of H2AK119-Ub IF (left) and H3K27me3 IF (right) of αCD3/αCD28 stimulated T cells treated with 2.5μM of IKKβ kinase inhibitors IMD-0354 or TPCA-1 for 48 hours. Averages of 3 independent experiments with SEM are shown with each dot representing a biological replicate. Statistical significance quantified using one way ANOVA followed by multiple comparisons to mock treated condition with 36-118 nuclei counted per condition per experiment. **F.** Volcano plot of differentially expressed X-linked genes comparing T cells stimulated in the absence or presence of IMD-0354, with genes escaping XCI in T cells stimulated in the presence of IMD-0354 colored in light blue. **G.** Volcano plot of genes differentially expressed from the Xi comparing T cells stimulated in the absence or presence of IMD-0354. **H.** Heatmap of differentially expressed (z score) *Xist* Interactome protein coding genes between unstimulated T cells (n=3), stimulated T cells (n=3), and stimulated T cells in the presence of 2.5μM IMD-0354 (n=3). Complete lists of upregulated and downregulated genes are available in Supplemental Table 4.

We next asked whether NF-κB-induced changes in *Xist* RNA localization is associated with impaired XCI maintenance. We performed allele-specific RNA-sequencing using T cells from female F1 mus x cast mice (**Figure 1A**) cultured with αCD3/αCD28 and the NF-κB inhibitor IMD-0354, as well as untreated stimulated T cells and unstimulated control cells. Principal component analyses (PCA) comparing treated to non-treated cells indicate that all replicates grouped together based on stimulation status and IMD-0354 treated cells were separate from untreated groups (**Supplemental Figure S4E**). We observed that NF-κB inhibition affects gene expression genome-wide (**Supplemental Figure S4F**), that the Xi is mostly dosage compensated (**Supplemental Figure S4G**) and that *Xist* RNA expression does not change with NF-κB inhibition (**Supplemental Figure S4H**). However, there are approximately 200 X-linked genes whose expression is altered with NF-κB inhibition (XCI escape genes in blue in **Figure 5F; Supplemental Table S3**). NF-κB inhibition specifically altered the allele-specific expression of some genes from the Xi (**Figure 5G**), including aberrant upregulation of *Tsix* which is typically not expressed in somatic cells including T cells.

Next, we asked whether NF-κB-mediated activation during T cell stimulation regulates the expression of *Xist* RNA binding proteins that are required to tether *Xist* RNA transcripts to the Xi. We compiled a list of 304 *Xist* RNA binding protein genes (the “*Xist* RNA Interactome”) previously identified using mouse and human somatic cells^42–45^, and identified which genes were differentially expressed when comparing expression levels in stimulated T cells (when *Xist* RNA is localized to the Xi) to unstimulated T cells or T cells stimulated in the presence of NF-κB inhibitor (172 out of 304 genes) (**Figure 5H and Supplementary Table S4**). We identified 43 *Xist* RNA Interactome genes (cluster 1) which are significantly downregulated in stimulated T cells treated with NFκB inhibitor and unstimulated T cells in comparison to stimulated T cells (**Figure 5H and Supplementary Table S4**). Cluster 1 includes *Trim6* and *Rad21* which have regulatory roles in XCI maintenance^46,47^ and are also downregulated in T cells from female mice with spontaneous lupus-like disease where *Xist* RNA localization is perturbed^48^. We also identified 27 *Xist* RNA Interactome genes (cluster 5) which are significantly upregulated in both NF-κB inhibited stimulated T cells and unstimulated T cells in comparison to stimulated T cells (**Figure 5H and Table S4**). This group of genes includes *Spen*, which functions in *Xist*-mediated gene repression during XCI initiation^49^ and *Rybp*, a member of the PRC complex which interacts with Yy1^50^, a protein required for *Xist* RNA localization to the Xi in lymphocytes^29,51^. Thus, the NF-κB pathway is required for Xist RNA retention at the Xi, and disruptions in this pathway impair the expression of some X-linked genes at the Xi and also the expression of Xist RNA interactome genes necessary for tethering, localization, and gene silencing.

### Conservation of NF-κB signaling for Xist RNA localization to the Xi in T cells from mice and humans

To further confirm that NF-κB signaling is necessary for stimulation-dependent localization of *Xist* RNA and heterochromatic marks to the Xi, we examined epigenetic features of the Xi in female mice with a genetic deletion which impairs NF-κB translocation into the nucleus. We generated mice with a T cell specific functional deletion of the IKKβ subunit that is specifically required to activate canonical NF-κB (**Figure 6A**). We mated IKKβ flox/flox mice, where loxP sites flank exon 3, with CD4-Cre recombinase mice to generate T cell specific IKKβ cKO mice (“IKKβ cKO”). Genotyping bands were of expected sizes (**Supplemental Figure S5B**) and similar to a previous report using an exon 3 IKKβ flox/flox mouse in the context of a B cell specific functional deletion^52^. We observed decreased IKKβ protein expression in T cells from IKKβ cKO mice, consistent with a deletion of the kinase domain-containing exon 3 (**Supplemental Figure S5C**). Animals were healthy with abundant splenic T cells (**Supplemental Figure S5A**), similar to previous observations using a different IKKβ flox/flox deletion of exons 6 and 7^53,54^. We isolated splenic CD3^+^ T cells from female IKKβ cKO/cKO, IKKβ cKO/+, and WT littermates, and stimulated them with αCD3/αCD28 for 48 hours before performing sequential *Xist* RNA FISH and H2AK119-Ub IF. Female IKKβ cKO/cKO T cells have significantly lower levels of *Xist* RNA accumulation at the Xi (**Figure 6B-C**). Stimulated IKKβ cKO/cKO T cells displayed H2AK119Ub and H3K27me3 foci at similar levels to WT T cells (**Figure 6B**, **Figure 6D**). As expected, splenic B cells from female IKKβ cKO/cKO and IKKβ cKO/+ (with wildtype levels of IKKβ) have *Xist* RNA ‘clouds’, reflecting T cell specific IKKβ deletion (**Supplemental Figure S5D**). Thus, genetic deletion of IKKβ significantly reduces *Xist* RNA localization at the Xi with minimal impact on H2AK119-Ub foci, corroborating the results of our studies using the chemical inhibitors of IKKβ.

**Figure 6.**
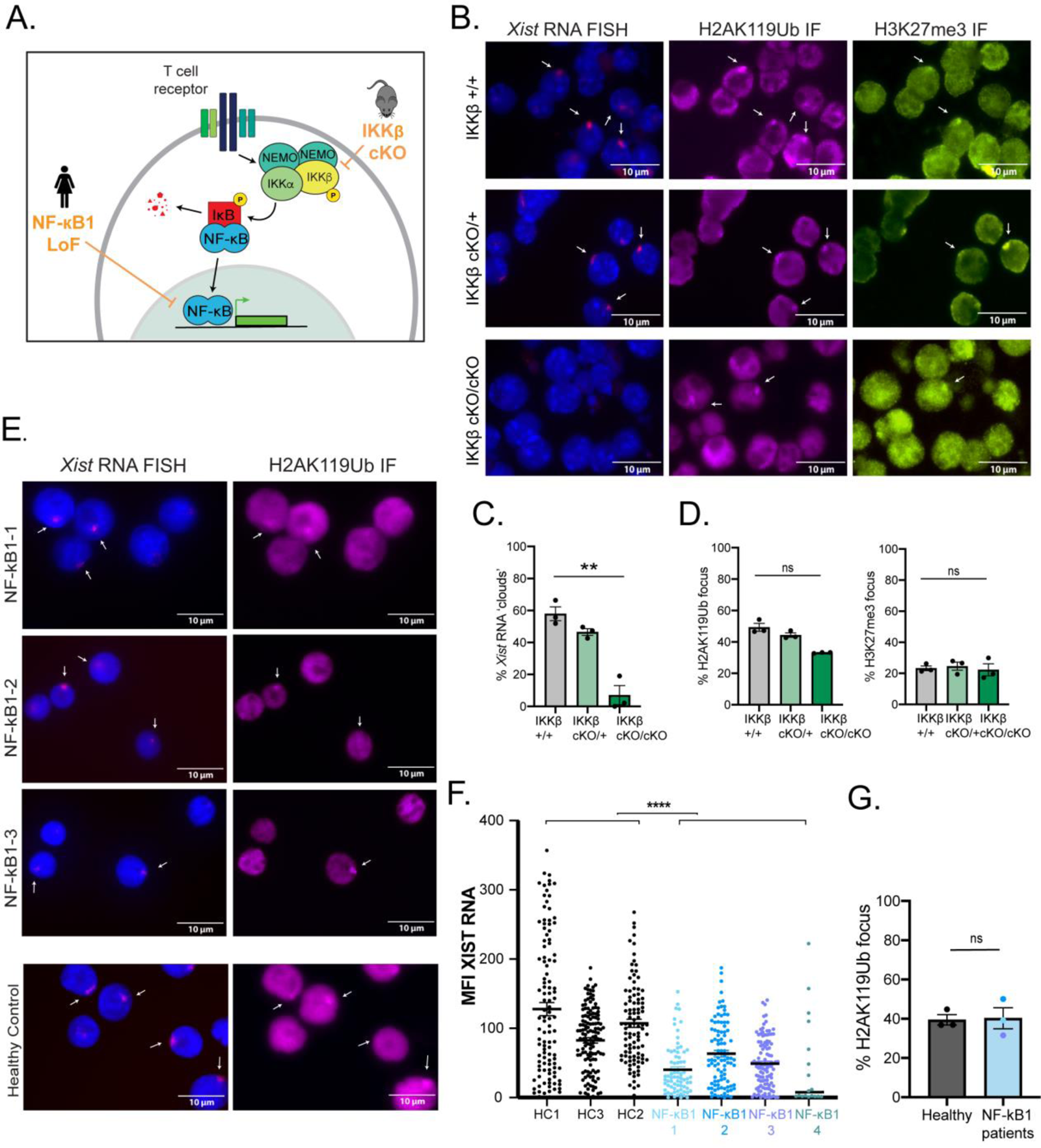
NF-κB signaling is required for Xist/XIST RNA localization to the Xi in activated T cells from mice and humans. **A**. Schematic of the loss of function mutations in mice and human heterozygous loss of function mutations. **B**. Representative fields (from 1 experiment) of *Xist* RNA FISH (left) H2AK119-Ub IF (middle) and H3K27me3 IF (right) of αCD3/αCD28 stimulated T cells from IKKβ +/+ (top), IKKβ cKO/+ (middle), and IKKβ cKO/cKO (bottom) female mice. **C**. Quantification of *Xist* RNA clouds of T cells from IKKβ +/+, IKKβ cKO/+, and IKKβ cKO/cKO female mice stimulated with αCD3/αCD28 for 48 hours. Averages of 3 independent experiments with SEM are shown with each dot representing a biological replicate. Statistical significance quantified using Kruskal-Wallis test, ** p<0.01, with 108-226 nuclei counted per condition per experiment. **D**. Quantification of H2AK119Ub foci (left) and H3K27me3 foci (right) of T cells from IKKβ +/+, IKKβ cKO/+, and IKKβ cKO/cKO female mice stimulated with αCD3/αCD28 for 48 hours. Averages of 3 independent experiments with SEM are shown with each dot representing a biological replicate. Statistical significance quantified using Kruskal-Wallis test, ** p<0.01, with 99-150 nuclei (H2AK119ub) or 58-122 nuclei (H3K27me3) counted per condition per experiment. **E.** Representative fields of *XIST* RNA FISH (left) and H2AK119-Ub IF (right) of sorted central/effector memory CD4^+^ T cells (CD45RA^-^CD27^+^) stimulated with αCD3/αCD28 for 48 hours from female NF-κB1 patients as well as a healthy control (healthy control #1). **F.** Mean fluorescence intensity (MFI) of *XIST* RNA for sorted central/effector memory CD4^+^ T cells from healthy controls (black) and NF-κB1 patients (light blue, blue, purple, teal) stimulated with αCD3/αCD28 for 48 hours, with each dot representing a nucleus. Statistical significance quantified using the Mann-Whitney U test comparing the MFI for XIST RNA (pooled HC1-3 vs. pooled NF-κB1 1-4, with 99-137 nuclei and 73-103 nuclei counted per HC and NF-κB1 sample, respectively), ****p<0.0001. Statistical significance remained the same when excluding Patient 4 (teal, 18 nuclei) because of disproportionately low cell numbers. **G**. Quantification of H2AK119Ub foci in sorted central/effector memory CD4^+^ T cells from healthy controls (black) and NF-κB1 patients (NF-κB1-1 teal, NF-κB1-2 blue, NF-κB1-3 purple) stimulated with αCD3/αCD28 for 48 hours. Averages with SEM are shown with each dot representing a patient (NF-κB1-1 teal, NF-κB1-2 blue, NF-κB1-3 purple). Statistical significance quantified using Kruskal-Wallis test, ** p<0.01, with 87-114 nuclei (healthy) or 36-76 nuclei (patients) counted per condition per experiment.

Finally, we asked whether the requirement for NF-κB signaling for *XIST* RNA localization and tethering to the Xi is conserved in human T cells. We collected peripheral blood mononuclear cells (PBMCs) from four female patients with heterozygous loss of function (LOF) mutations in the NF-κB1 protein (p50; labeled NF-κB1-1, 2, 3, 4), which is one of the subunits of canonical NF-κB (**Supplemental Table S5** and **Figure 6A**). The specific NF-κB1 (p50) LOF mutations in these patients include either a nonsense mutation (NFκB1-1) or two frameshift mutations (NF-κB1-2, NF-κB1-3, NF-κB1-4)^55,56^ (**Supplemental Table S5**), and are all associated with the clinical phenotype of Common Variable Immune Deficiency (CVID), an immunodeficiency that is often complicated by autoimmune phenotypes^57^. The NF-κB1 protein (p50) contains a RHD domain but lacks a transactivation domain, and p50 proteins only drive transcription when bound to transactivation domain containing NF-κB proteins including p65 (RelA)^58^. Moreover, while the prototypic canonical p65:p50 heterodimer promotes transcription, p50:p50 homodimers act as transcriptional repressors^59,60^ Thus, in contrast to IKKβ deletions, NF-κB1 loss of function mutations can have diverse impacts on downstream gene activation and repression. We sorted central/effector memory CD4^+^ (CD45RA^-^CD27^+^) and, due to lower cell quantities, bulk CD8^+^ T cells from NF-κB1-1, 2, 3, 4 patients and age-matched healthy controls (**Supplemental Figure S5E**), stimulated cells *in vitro* with αCD3/αCD28 for 2 days, and subsequently performed sequential XIST RNA FISH and IF. We detected XIST RNA signals in heterozygous null NF-κB1 patients, yet we observed that the intensity and size of the XIST RNA ‘clouds’ was significantly smaller with reduced fluorescence intensity compared to healthy controls (**Figure 6E**). Quantification of the mean fluorescence intensity (MFI) of the nuclear XIST RNA signals for *in vitro*-stimulated T cells from NF-κB1-1-4 patients and healthy controls confirmed that NF-κB1 heterozygous null mutation significantly reduces XIST RNA ‘clouds’ at the Xi (**Figure 6F** and **Supplemental Figure 5F**). We observed similar levels of H2AK119-Ub foci in central/effector memory CD4^+^ T cells from heterozygous NF-κB1 null patients compared to healthy controls (**Figure 6G**), indicating that reduced XIST RNA accumulation at the Xi does not affect enrichment of H2AK119-Ub. Together these results demonstrate that NF-κB signaling regulates *Xist/XIST* RNA localization at the Xi in activated T cells from both mice and humans.

## DISCUSSION

T cells are a unique example of somatic cells where epigenetic modifications are dynamically recruited to the Xi upon cellular activation. In this study, we investigated the transcriptional status and heterochromatic histone mark enrichment of the Xi in unstimulated T cells, and how gene expression and epigenetic modifications change across the Xi with *in vitro* stimulation. We discovered that unstimulated T cells lacking cytological enrichment of Xist RNA and heterochromatic marks H3K27me3 and H2AK119-Ub are dosage compensated, and that there is detectable enrichment of H3K27me3 modifications across the Xi at expressed and silent genes. T cell stimulation with anti-CD3/CD28 results in accumulation of H2AK119-Ub modifications across the Xi, and increased accumulation of H3K27me3. Furthermore, we dissected the requirements for the dynamic relocalization of Xist RNA and H2AK119-Ub to the Xi upon T cell activation by examining specific components of the T cell receptor signal transduction cascade. To our knowledge, these data are a novel link between NF-κB signaling, dynamic localization of XIST/Xist RNA to the Xi and regulation of expression of some X-linked genes from the Xi in female immune cells.

The Xi has higher levels of H3K27me3 modifications compared to the Xa in unstimulated T cells, yet levels are not high enough to be visualized cytologically. Using allele-specific CUT&RUN, we achieve greater sensitivity and resolution to detect specific locations across the Xi with enrichment of H3K27me3 and H2AK119-Ub. There are similar enrichment levels for H2AK119-Ub across the Xi and Xa in unstimulated T cells, which aligns with our predictions for both H2AK119-Ub and H3K27me3 based on our previous cytological analyses^27,29^. We were surprised to see that both H3K27me3 and H2AK119-Ub are enriched at the upstream promoter regions of XCI escape genes compared to repressed genes (**Figure 1E**), as these modifications are typically associated with gene repression^61,62^. T cells are dosage compensated, yet there is significant amount of transcription from the Xi in both unstimulated and stimulated cells. Our analyses show that there are ∼55 X-linked genes that escape XCI in T cells (**Figure 2**), and about 20 X-linked genes are specifically expressed from the Xi in unstimulated T cells which lack *Xist* RNA localization to the Xi^27^. Both H3K27me3 and H2AK119-Ub are associated with facultative chromatin, which allows for gene reactivation, and the Xi is considered an example of facultative heterochromatin^63^. H3K27me3 enrichment across the gene body is the typical pattern for transcription inhibition, which is observed for repressed genes on the Xi (black line, **Figure 1E**). However, there are transcribed genes with H3K27me3 enrichment at promoter regions for various cell types in mice^64^. The role of enrichment of H3K27me3 at promoters of XCI escape genes is unknown, but it may have a repressive function to moderate gene expression levels from the Xi. Unstimulated T cells are quiescent, and there are reduced global levels of H3K27me3 in quiescent lymphocytes,^65^ yet the Xi retains enrichment of this modification and not H2AK119-Ub. We propose that H3K27me3 enrichment across the Xi may function as an epigenetic imprint to direct Polycomb Repressive Complex 1 (PRC1)-mediated accumulation of H2AK119-Ub following T cell stimulation through the TCR. It is well established that H3K27me3 modifications can recruit canonical PRC1 recruitment, through interactions with CBX family proteins containing a chromodomain^66–68^, and it remains to be determined whether a canonical PRC1 recruitment pathway functions in stimulated T cells. During the formation of the Xi, H2AK119-Ub modifications accumulate first on the Xi through deposition by non-canonical PRC1 complex and precede PRC2-mediated H3K27me3 accumulation^69^. This sequence of events is in stark contrast to the order of histone modifications observed during T cell stimulation. Additional experiments are required to determine if H3K27me3 modifications function as an epigenetic imprint for PRC1 recruitment and H2AK119-Ub deposition across the Xi in unstimulated T cells.

TCR engagement is necessary for both *Xist* RNA and H2AK119-Ub localization to the Xi when T cells are activated. Curiously, the TCR peptide affinity does not affect the percentages of nuclei with *Xist* RNA and H2K119-Ub localization at the Xi beyond a minimal signaling threshold, as we did not observe graded re-localization of *Xist* RNA and H2K119-Ub with intermediate affinity peptides SII<u>V</u>FEKL and SII<u>Q</u>FEKL, but instead saw minimal re-localization with the low affinity peptide <u>E</u>IINFEKL (**Figure 3**). Some studies using a different intermediate affinity peptide (SII<u>T</u>FEKL) have demonstrated sub-maximal NF-κB activation compared to the high affinity SII<u>N</u>FEKL peptide^70,71^. It is not clear whether NF-κB activation occurs after stimulation with the low affinity <u>E</u>IINFEKL peptide. Yet, our data suggests NF-κB signaling, regardless of whether it reaches maximal levels, is sufficient for *Xist* RNA re-localization in T cells. It is possible that that IL-2 signaling, independent of TCR engagement, also functions to activate gene expression pathways necessary for Xist RNA and H2K119-Ub enrichment at the Xi when T cells are stimulated. Our study found that IL-2 alone results in Xist RNA ‘clouds’ in ∼5% of nuclei, yet CD28 receptor activation alone is does not localize either Xist RNA or H2K119-ub to the Xi (**Figure 4C**).

Xist RNA localization to the Xi, and expression of some X-linked genes, requires NF-κB signaling activity in T cells. With chemical or genetic disruption of IKKβ, stimulated T cells have visible H2AK119-Ub and H3K27me3 foci that co-localize with an X chromosome, despite lack of localized *Xist* RNA. PRC1 and PRC2 are responsible for deposition of H2AK119-Ub and H3K27me3, respectively, and NF-κB mediated inhibition of Xist RNA localization at the Xi suggests that during XCI maintenance, PRC1 and PRC2 are recruited to the Xi independently of *Xist* RNA in T cells. The epigenetic imprint responsible for this recruitment could be H3K27me3 modifications on the Xi, which are abundant across the Xi in unstimulated T cells (**Figure 1D**). Culturing T cells with IKKβ kinase inhibitors prevented Xist RNA accumulation at the Xi and altered the expression of ∼200 X-linked genes, and specifically 14 genes on the Xi (**Figure 5F-G**). We did not observe differences in *Xist* gene expression with NF-κB inhibition, indicating that *Xist* is not a target of NF-κB mediated repression or activation. The observed reduction of Xist RNA ‘clouds’ is therefore not due to NF-κB-mediated transcriptional effects on the *Xist* gene, but is instead likely the result of altered expression of Xist RNA Interactome protein genes which are required for tethering and localizing Xist RNA transcripts to the Xi (**Figure 5H**). There are 43 Xist Interactome genes in cluster 1 that are downregulated in NF-κB inhibited T cells, and additional work is needed to determine how these factors anchor Xist RNA transcripts to the Xi specifically in activated T cells and how NF-κB regulates their expression. There are many NF-κB target genes encompassing a variety of cellular functions that have been described (https://www.bu.edu/nf-kb/gene-resources/target-genes/), some of which are X-linked, and our results using NF-κB inhibition suggest that there are many X-linked genes whose expression is influenced by NF-κB. Our study suggests that XCI maintenance and Xist RNA Interactome genes are a new category for NF-κB mediated regulation that should be considered.

Our data suggest a novel pathway for understanding why patients with TCR and B cell receptor signaling defects can have immunodeficiency and also features of autoimmunity. It has been traditionally thought that mutations in TCR and BCR proteins would favor selection of autoreactive receptors. Our findings that NF-κB inhibition alters X-linked gene expression on both the Xa and Xi suggests that loss of TCR signaling can promote altered expression of pro-inflammatory genes in female patients with defects in TCR signaling. It is possible that altered X-linked expression might also be a feature in female patients with mutations in the IKK complex, for example IKBKG-deficiency (incontinentia pigmenti), subtypes of which have been associated with inflammatory complications^72,73^. Additional investigation of X-linked expression profiles in these patient populations is necessary to determine whether there is a role for altered XCI maintenance in these diseases. We have previously reported aberrant patterns of *XIST/Xist* RNA localization in T and B cells from SLE patients and lymphocytes from spontaneous mouse models of lupus-like disease ^27,48,74^. SLE patient T cells can exhibit abnormal NF-κB activity^75^ and NF-κB is an important modulator of autoimmune disease progression. Furthermore, NF-κB activity in T cells is critical for protection during immune challenge, thus understanding the molecular mechanisms between NF-κB signaling and the regulation of XCI maintenance will shed light on understanding the genetic and epigenetic contributions underlying sex-biased immune responses during infection.

## MATERIALS AND METHODS

### Ethics statement

Animal studies performed in this study fall under an animal protocol approved by the University of Pennsylvania Institutional Animal Care and Use Committee (protocol # 804864). The University of Pennsylvania is an AAALAC accredited institution and adheres to the standards set by the Animal Welfare Act and the NIH Guide for the Care and Use of Laboratory Animals.

### Mouse Models

C57Bl/6j (strain# 000664, Jackson) mice were bred in our colony and were used unless otherwise indicated. F1 mice were bred in our colony in the following manner: Xist^fl/fl^ female mice were a gift from R.Jaenisch^76^ and backcrossed to C57BL/6j (strain# 000664, Jackson) for 10 generations, then mated to an ACTB-Cre male (B6N.FVB-Tmem163Tg(ACTB-cre)2Mrt/CjDswJ; strain# 019099, Jackson) to generate heterozygous Xist^fl/+^ females. Heterozygous females were then bred with wild-type Mus castaneus (Cast) males to generate F1 Xist^fl/+^ female mice. F1 Xist^fl/+^ females always inactivate the paternal WT Cast X chromosome. Spleens from naïve OT-I mice (strain# 003831, Jackson) and CD45.1 C57Bl/6 mice (strain# 002014, Jackson) were a gift from Christopher Hunter (University of Pennsylvania). IKKβ cKO mice were bred in our colony by crossing CD4cre mice (strain# 022071, Jackson) to IKKβ flox/flox mice^52^ (a gift from Michael May, University of Pennsylvania). For genotyping the IKKβ flox locus the following primer sequences were used: 5’-CAC AGT GCC CAC ATT ATT TAG ATA GG -3’ and 5’-GTC TTC AAC CTC CCA AGC CTT-3’, with expected amplification sizes of ∼200bp (flox allele) and ∼180bp (WT allele). All mice utilized for this study were females ranging from 8-14 weeks in age. All animals were age matched within an individual experiment. Euthanasia via carbon dioxide followed by cervical dislocation was used for animal sacrifice prior to isolations. All mice were maintained at the Penn Vet animal facility, and experiments were approved by the University of Pennsylvania Institutional Animal Care and Use Committee (IACUC).

### Murine T cell isolation, stimulation, cell cycle analyses and NF-κB inhibition

Spleens were ground using glass slides prior to passage through a 70uM filter and red blood cells were lysed with ACK lysis buffer (Quality Biological). Murine splenic CD3^+^ T cells were isolated using CD3^+^ T cell enrichment columns (R&D Systems, MTCC25). T cells were cultured at 37C in RPMI-1640 (Invitrogen) containing 10% FBS (Gemini), 0.1% β-mercaptoethanol (Invitrogen), 1% nonessential amino acids (Invitrogen), 1% sodium pyruvate (Invitrogen), and 1% penicillin-streptomycin (Invitrogen) (T cell media). T cells were stimulated with various combinations of stimulating reagents (indicated in the figure legends): 1 μg/ml plate-bound αCD3 (Bio X cell, BE0001-1-5mg), 10 μg/ml soluble αCD28 (Bio X cell, clone 37.51), 20 U/ml soluble recombinant IL-2 (Biolegend) and cultured for 48-72 hours. Cells were harvested for slide preparation at 48 or 72 hours (as indicated in each figure).

For cell cycle analysis, CD3^+^ T cells were cultured in the presence of αCD3 and αCD28 similarly to previous studies^35^. Briefly, either 200μM hydroxyurea (S phase), 1μg/ml nocodazole (G2/M), or a mock treatment control was added to the culture medium. For rapamycin treatment, cells were re-suspended in T cells media containing 100nM rapamycin and incubated at 37C for 4 hours prior to culturing in the presence of αCD3/αCD28.

For calcium analysis, CD3^+^ T cells were cultured in DMEM (Invitrogen) or calcium-free DMEM (Gibco) containing 10% FBS (Gemini), 0.1% β-mercaptoethanol (Invitrogen), 1% nonessential amino acids (Invitrogen), 1% sodium pyruvate (Invitrogen), and 1% penicillin-streptomycin (Invitrogen). For NF-κB chemical inhibition, CD3^+^ T cells were cultured in the presence of αCD3 and αCD28 as well as with 2.5μM, 1μM, or 0.5μM IMD-0354 (Cayman Chemical) or TPCA-1 (Cayman Chemical).

For isolation of splenic T cells from IKKβ cKO mice, B cells were first isolated using positive CD23^+^ enrichment (see “B cell isolation and culturing” below) and the flowthrough was then treated to red blood cell lysis and CD3^+^ T cell enrichment as above.

### MEF culturing

Mouse Embryonic Fibroblasts (MEF) were cultured in DMEM supplemented with 10% FBS (Gemini) and 1% penicillin-streptomycin (Invitrogen). Cells were seeded onto six well plates and when 80% confluent the media was supplemented with 10μM, 5μM, 2.5μM, 1μM, or 0.5μM IMD-0354 (Cayman Chemical) or TPCA-1 (Cayman Chemical).

### Allele-specific RNA sequencing

T cells from F1 mus x cast female mice at either 0 hour (n=6), 48 hours post stimulation (n=6), or 48 hours post stimulation in the presence of 2.5μM IMD0354 (n=3) were collected into TRIzol reagent (15596026, ThermoFisher). RNA isolations were performed according to the manufacturers protocol. Libraries were prepared with an Illumina TruSeq Stranded Total RNA Stranded Ligation Ribo-Zero+ kit (20040529, Illumina), pooled, and run on an Illumina NextSeq 2000 sequencer (150bp paired-end). For determining XCI escape genes, we performed experiments using n = 3 mice, and then utilized another set of n = 3 female mice for NF-κB inhibition studies.

To quantify F1 T cell gene expression allele-specifically, we created an N-masked mm10 (C57BL/6j; Ensembl GRCm38) genome using SNPsplit^77^. SNPs were derived from the Casteneus genome for the C57BL6 x Cast F1 mice. The resulting N-masked genomes were used to generate respective STAR (v2.7.1a) indexes. Reads were aligned using STAR with alignEndsType set to EndToEnd, and outSAMattributes set to NH HI NM MD to allow for compatibility with SNPsplit. STAR-aligned files were then passed through SNPsplit for allele-specific sorting of reads. Reads were quantified using featureCounts^78^ on both the diploid N-masked aligned and the haploid allele-specific aligned reads.

Genes that escape XCI were identified using 3 thresholds of escape, as previously described ^21,28^. Briefly, diploid gene expression was first calculated in RPKM (reads per kb of exon length, per million mapped reads), and genes were called as expressed if their diploid RPKM was > 1. For every X-linked gene that passed this threshold, haploid gene expression was calculated in SRPM (allele-specific SNP-containing exonic reads per 10 million uniquely mapped reads), and genes which had an Xi-SRPM > 2 were considered to be expressed from the Xi. Finally, a binomial model estimating the statistical confidence of XCI escape probability was applied to the genes passing the first 2 thresholds. This model compares the proportion of Xi-specific reads to the total Xi + Xa reads and calculates a 95% confidence interval. If the 95% lower confidence limit of a gene’s escape probability was greater than 0, it was called a gene escaping XCI. Our code for calling XCI escape genes (escape_gene_calculator) is publicly available on github (https://github.com/Montserrat-Anguera/escapegenecalculator). Genes that escaped XCI were grouped by their escape status in unstimulated and stimulated cells, generating 3 different categories: genes that escape in unstimulated cells only, genes that escape in stimulated cells only, and genes that escape in both unstimulated and stimulated cells.

Mapped reads were graphed using Prism v8.4.3. Statistical significance of total RPM (X) was determined using unpaired T test with Welch’s correction. To graph the log SRPM of Xa vs Xi transcripts an arbitrary value of 0.01 was added to all SRPM values. To calculate significance, the SRPM of Xi divided by SRPM (Xi+Xa) was calculated for each gene in the category of either XCI escape or XCI subject genes and then analyzed with a Mann-Whitney non-parametric T test. Venn diagram of escape genes was generated using Biovenn^79^.

For differential gene analysis between unstimulated, stimulated, and stimulated + NF-κB inhibition conditions, we used the N-masked index created above and aligned using STAR with alignEndsType set to SortedyCoordinate, and quantmode set to GeneCounts. STAR-aligned files were then quantified using featureCounts^78^. Reads were filtered to have a Counts per million (CPM) >1 for at least 3 samples prior to quantile normalization. For PCA analysis, distance was calculated using the dist function (Method= “Euclidean”) and clusters were calculated using the hclust function (Method = “complete”). Differetial gene lists were generated using the DeSeq2 package^80^. For differential gene expression from the Xi, the above steps were used with the Castaneus read file generated from SNP-split. For *Xist* RNA interactome analysis, a list of known binding proteins was compiled manually from published literature^42–45^. This list was filtered to exclude genes that were not found to be differentially expressed in either unstimulated T cells vs stimulated T cells or stimulated T cells vs T cells stimulated with 2.5μM IMD0354. Heatmaps of the remaining 172/304 genes were generated using hclust function (Method = “spearman”) and plotted using the heatmap2 function.

### CUT&RUN for allele-specific epigenomic profiling of Xa and Xi

CUT&RUN for H3K27me3 and H2AK119Ub was performed as previously described^81^. In brief, 10uL/reaction of concanavalin A-coated beads (Cat #86057-3, Polysciences) were equilibrated with Binding Buffer (20mM HEPES, 10mM KCl, 1mM CaCl2, 1mM MnCl2) then resuspended in Wash Buffer (20mM HEPES, 150mM NaCl, 0.5mM Spermidine, 1x PIC (Cat #11697498001, Roche)). Approximately 1 million naïve or 0.7 million stimulated CD3^+^ T cells were immobilized on beads in Wash Buffer by rotating for 1 hr at room temperature. Beads were resuspended in Antibody Buffer (20mM HEPES, 150mM NaCl, 0.5mM Spermidine, 1x PIC, 0.05% Digitonin, 2mM EDTA). 3ug of primary antibody for H3K27me3 (Cat #39055, Active Motif), H2AK119Ub (Cat # 8240S, Cell Signaling), or IgG (Cat #A01008, GenScript) were added and beads were rotated at 4°C for 5 hr. Beads were washed with and resuspended in Dig-Wash Buffer (20mM HEPES, 150mM NaCl, 0.5mM Spermidine, 1x PIC, 0.05% Digitonin). 1200ng/mL of pA-MNase fusion protein (gift from Henikoff Lab) was added and beads were rotated at 4°C for 1 hr. Beads were washed twice then resuspended in Dig-Wash Buffer. Targeted digestion was performed at 0°C for 30 min by adding CaCl2 to 2mM. The digestion was stopped by adding one volume of 2X STOP Buffer (340mM NaCl, 20mM EDTA, 4mM EGTA, 0.02% Digitonin, 50ug/mL RNase A (Cat #3335399001, Roche), 50ug/mL Linear Acrylamide (Cat #K548, VWR), 2pg/mL Yeast Spike-In DNA). The target chromatin was released into the supernatant by shaking beads at 300rpm for 10 min at 37°C. The supernatant was then transferred to fresh tubes and incubated at 70°C for 10 min with SDS to 0.1% and 50ug of Proteinase K (Cat # EO0491, Thermo Scientific). DNA was then isolated with phenol:chloroform:isoamyl alcohol followed by precipitation with ammonium acetate. Libraries were prepared with 5ng of CUT&RUN DNA using NEBNext Ultra II DNA Library Prep Kit for Illumina (Cat # E7645S, NEB) following manufacturers protocol. CUT&RUN libraries were then paired-end sequenced on a P3 300 flow cell on the NextSeq2000.

Data Processing for Allele-Specific CUT&RUN was performed as follows. Adapters were removed using Trimmomatic (V0.32, https://github.com/usadellab/Trimmomatic). SNPsplit^77^ (0.5.0) was used to generate an “N-masked” version of the mouse reference genome (mm10), wherein all SNPs between the Mus musculus C57BL/6 and Mus musculus CAST/EiJ mouse strains are masked by an “N” nucleotide. Reads were mapped to the “N-masked” genome using Bowtie2 (2.3.4.1) with the parameters [-q -- local --very-sensitive-local --soft-clipped-unmapped-tlen --dovetail --no-mixed --no-discordant --phred33 -I 10 -X 1000]. Duplicated reads were removed using Picard MarkDuplicates (1.141, https://broadinstitute.github.io/picard/) with option [REMOVE-DUPLICATES=true]. Reads mapped to regions from the ENCODE blacklist were removed. Allele-specific BAM files were then generated using SNPsplit (0.5.0), wherein reads overlapping SNPs were tagged and sorted into Bl6 and Cast files. CPM normalized bigwig files were generated using deepTools bamCoverage (3.5.1, https://deeptools.readthedocs.io/en/develop/) with options [--effectiveGenomeSize 2652783500 --normalizeUsing CPM --ignoreForNormalization chrX]. CPM normalized biological replicates were combined with WiggleTools mean (1.0, https://github.com/Ensembl/WiggleTools) and visualized with IGV.

Heterochromatic mark density across gene bodies was determined using EnrichedHeatmap^82^ (1.22.0), which plotted heterochromatic mark density ±5kb of X-linked genes expressed in T cells. X-linked genes expressed in T cells were defined as those with an RPKM >1 in ≥2 biological replicates in our allele-specific RNAseq analysis. Expressed genes were partitioned by escape status as determined with the previously discussed cutoffs.

D-Score analysis was performed by defining promoter regions as ±1kb from the TSS for all X-linked genes. Gene bodies were defined 1kb downstream of the TSS to the TES. Gene bodies of X-linked genes shorter than 1kb were excluded from the analysis. Intergenic regions were defined as 10kb windows (generated using GenomicRanges (1.44.0)) that did not overlap gene bodies or their promoter regions (1kb upstream of TSS). Reads were assigned to defined windows using featureCounts from the Rsubread package (2.6.4) with arguments [useMetaFeatures = F, isPairedEnd = T]. For each biological replicate, d-scores [(readsB6/(readsCast+ readsB6))-0.5] were calculated for promoters, gene bodies, and intergenic regions. Biological replicates were then averaged for each timepoint. P-values were calculated using a Wilcoxon signed-rank test with Benjamini Hochberg correction.

### Sequential Xist RNA FISH and H2AK119Ub, H3K27me3 Immunofluorescence, & MFI quantification of XIST RNA

Sequential RNA FISH and immunofluorescence (IF) for murine CD3^+^ T cells was performed following established protocols^26,27^, where *Xist* RNA FISH was performed first, followed by histone modification IF. Briefly, cells were cytospun onto glass slides, permeabilized with CSK-T buffer (100mM NaCl, 300mM sucrose, 10mM PIPES, 3mM MgCl_2_, 0.5% triton, pH6.8), fixed with 4% paraformaldehyde for 10 minutes, and dehydrated with a series of increasing ethanol concentrations. For *Xist* RNA FISH, 2 Cy3-labeled 20-nucleotide oligo probes were designed to recognize regions within exon 1 (synthesized by IDT). For IF, cells were blocked with 0.2% PBS-Tween, 0.5% BSA. Histone H2AK119-Ub (Cell Signaling; catalog # 8240S) and Histone H3K27me3 (Active Motif; catalog# 39155) antibodies were diluted 1:100.

Sequential *XIST* RNA FISH and H2AK119-Ub IF for sorted and stimulated human T cell populations was performed following established protocols. For *XIST* RNA FISH, 3 Cy3-labeled 20-nucleotide oligo probes recognizing repeats across *XIST* RNA were synthesized (IDT)^74^ . After imaging FISH, we performed IF. Cells were blocked with 0.2% PBS-Tween, 5% BSA. Histone H2AK119-Ub (Cell Signaling; catalog # 8240S) and Histone H3K27me3 (Active Motif; catalog# 39155) antibodies were diluted 1:100. Images were obtained using a Nikon Eclipse microscope and murine samples were initially categorized by the 4 types of *Xist* RNA localization patterns, as described previously^26,27,83^. Nuclei were called as displaying localized *Xist* RNA when they were categorized as type I or II (pinpoints of *Xist* RNA visible in a localized area of the nucleus the size of one X). Statistical significance was calculated using unpaired T tests with Welch’s correction and one-way ANOVA followed by multiple comparisons as indicated in figure legends.

Mean fluorescence intensity (MFI) analysis for *XIST* RNA in human T cells was performed using Fiji^84^ (2.9.0/1.53t). Briefly, DAPI and *XIST* (Cy3) channels were imported separately into Fiji as greyscale images and a Z-projection (MAX) image of each was generated using in focus Z-stacks. The DAPI image was then threshold adjusted (default setting), made binary, and then watershed corrected to account for nuclei touching. The corrected DAPI image was then used to generate regions of interest (ROI), with size = 0.3 micron-infinity and circularity = 0.5-1. Any burst nuclei or debris that passed these thresholds were manually excluded following comparison to the original image. The Cy3 image was background corrected in the following manner: the MFI of a rectangle 150 pixels x 100 pixels outside of the area of any ROIs or debris was calculated using the measure command within the ROI manager, and the resulting MFI was subtracted from the image. The ROIs generated from the corrected DAPI image were overlaid on to the background corrected Cy3 image and the MFI measured for each ROI in the image.

### OT-I experiments & cell sorting

Splenocytes were isolated from the spleen of a C57Bl/6 (CD45.1^+^) mouse, and peptide pulsed in the following manner. Briefly, splenocytes were incubated with 1uM of SIINFEKL, SIIQFEKL, SIIVFEKL, EIINFEKL, or mock for 20 minutes. Peptides were a gift from Christopher Hunter (University of Pennsylvania) and synthesized by Biosyn^32^. CD3^+^ T cells were isolated from the spleens of female OT-I (CD45.1^+^/CD45.2^+^) mice using CD3^+^ T cell enrichment columns (R&D Systems, MTCC25) and co-cultured with the peptide-pulsed splenocytes for 3 days. Following Fc receptor blockade with anti-mouse CD16/CD32, cells were stained with anti-CD45.1, anti-CD45.2, and anti-CD8α. CD45.1^+^/CD45.2^+^/CD8α^+^ OT-I T cells were sorted on a FACS Jazz sorter (Children’s Hospital of Philadelphia) and slides were immediately prepared for RNA FISH.

### Human NF-κB1 patient samples and sorting

Human peripheral blood mononuclear cells (PBMCs) from deidentified healthy female individuals were collected by the University of Pennsylvania Pathology BioResource Human Immunology Core facility. The average age of the healthy control group was 54.6 years (range=52-58 years old). Human PBMCs from deidentified female patients with mutations in NF-κB1 were collected from: the Children’s Hospital of Philadelphia (NF-κB1-1); the Icahn School of Medicine at Mt. Sinai (NFκB1-2), Massachusetts General Hospital (NF-κB1-3); and the Medical College of Wisconsin (NF-κB1-4). The average age of the patient group was 59.5 years (range=52-75 years old). PBMCs were thawed and staining for surface antigens was performed following Fc receptor blockade with Human TruStain FcX (BioLegend). Central/effector memory CD4^+^ T cells were defined as live/CD16^-^/CD14^-^/CD123^-^/CD56^-^/CD11c^-^/CD19^-^/CD4^+^ > excluding CD25^hi^CD127^lo^ > CD45RA^-^/CD27^+^. CD8^+^ T cells were defined as live/CD16^-^/CD14^-^/CD123^-^/CD56^-^/CD11c^-^/CD19^-^/CD4^-^CD8^+^. Due to low cell numbers, CD8^+^T cells from patient NF-κB1-4 were not isolated. T cells were sorted using a 5-laser Cytek Aurora CS (Children’s Hospital of Philadelphia). Sorted T cells were stimulated with Human T-Activator Dynabeads (Thermofisher) in RPMI-1640 (Invitrogen) containing 10% FBS (Gemini), 1% sodium pyruvate (Gibco), 1% GlutaMax (Gibco), and 1% penicillin-streptomycin (Invitrogen) for 2 days before cells were harvested and slides prepared. Because we examined human T cells for *XIST* RNA cloud and repressive histone modifications localization on the Xi, we only used female patient and healthy control samples.

### Western blot

T cells from IKKβ cKO/cKO, IKKβ cKO/+ or WT littermates were isolated and cell lysates were made from either unstimulated or stimulated T cells using 1X RIPA buffer plus protease inhibitor. Lysates were sonicated for 3 pulses using a probe sonicator with dial set to 5. 9ug of cell lysates were loaded onto a 4-12% Bis-Tris NuPage gel (Invitrogen) and run using MOPS running buffer. The gel was transferred using a wet transfer system and 10% methanol in Bis-Tris transfer buffer. Blot was cut and blocked for 1 hour at room temperature in 5% dehydrated milk in TBS-T. Blots were then incubated overnight at 4C with either anti-IKKβ (Cell Signaling, clone D30C6) or anti-β actin (Cell Signaling, clone 8H10D10) in 5% BSA TBS-T. Blots were then washed prior to incubation for 1 hour at room temperature with either rabbit (Cell Signaling, catalog # 7074S) or mouse (Cell Signaling, catalog # 7076S) HRP-conjugated secondary antibodies in 5% BSA TBS-T. Blots were developed using Pierce ECL substrate (ThermoFisher).

## Code availability

Code for escape gene analysis can be found on GitHub: https://github.com/Montserrat-Anguera/escapegenecalculator

## Data availability

All sequencing data generated in this study has been deposited to the NCBI GEO database.

## ACKNOWLEDGMENTS

We thank A. Ager and members of the Anguera, May, and Hunter labs for helpful discussions. We also thank the PennVet CHMI for their help with RNA and CUT&RUN sequencing, and J. Murray and the CHOP flowcore for help with cell sorting. We thank H. Wang for streamlining the allele-specific RNAseq code and generating the GitHub Escape_Gene_Calculator code, S. Henikoff for pA-MNase used for CUT&RUN experiments, and B. Freedman for sharing calcium deficient media. C. Lengner, W. Tong, and A. Kashina labs for sharing reagents.; M. Karin for providing the IKKβ flox/flox mice, and C. Hunter for providing spleens from OT-I and congenic WT mice. This research was supported by Lupus Research Alliance (Target in Lupus), NIH R01 AI134834, AI168047 to MCA; NIH R01 AI146026 and Jeffrey Model Foundation to N.R.; NIH T32 DK-07780 (to K.S.F.), NIH R01 AR066567, NIH R21 AI173679 (to M.J.M), NIH T32-AR076951-01 (to N.J.), H. Ralph Schumacher Rheumatology Research Fund (to N.J.), the Benjamin and Mary Siddons Measey Foundation (to N.J.), and the Scleroderma Research Foundation (to N.J.); SB is supported by the National Institute of Allergy and Infectious Diseases of the National Institutes of Health under Award Number K23AI163350.

## AUTHOR CONTRIBUTIONS

Conceptualization: K.S.F., N.E.T., M.J.M., N.R., M.C.A.

Methodology: K.S.F., N.E.T., N.J., L.S., M.C.A.

Investigation: K.S.F., N.E.T., A.D., N.J.

Formal Analysis: K.S.F., N.E.T., A.D.

Resources: L.S., C. C-R., S. B., J. R., N. R., M.J.M., M.C.A.

Manuscript-original draft: K.S.F. and M.C.A.

Manuscript Review and Editing: K.S.F., N.E.T., A.D., N. J., L. S., N.R., M.J.M, M.C.A.

Funding Acquisition: K.S.F. and M.C.A.

