## Supplemental Figures for "NF-κB Signaling is Required for X-Chromosome Inactivation Maintenance Following T cell Activation"

**SUPPLEMENTAL MATERIAL**

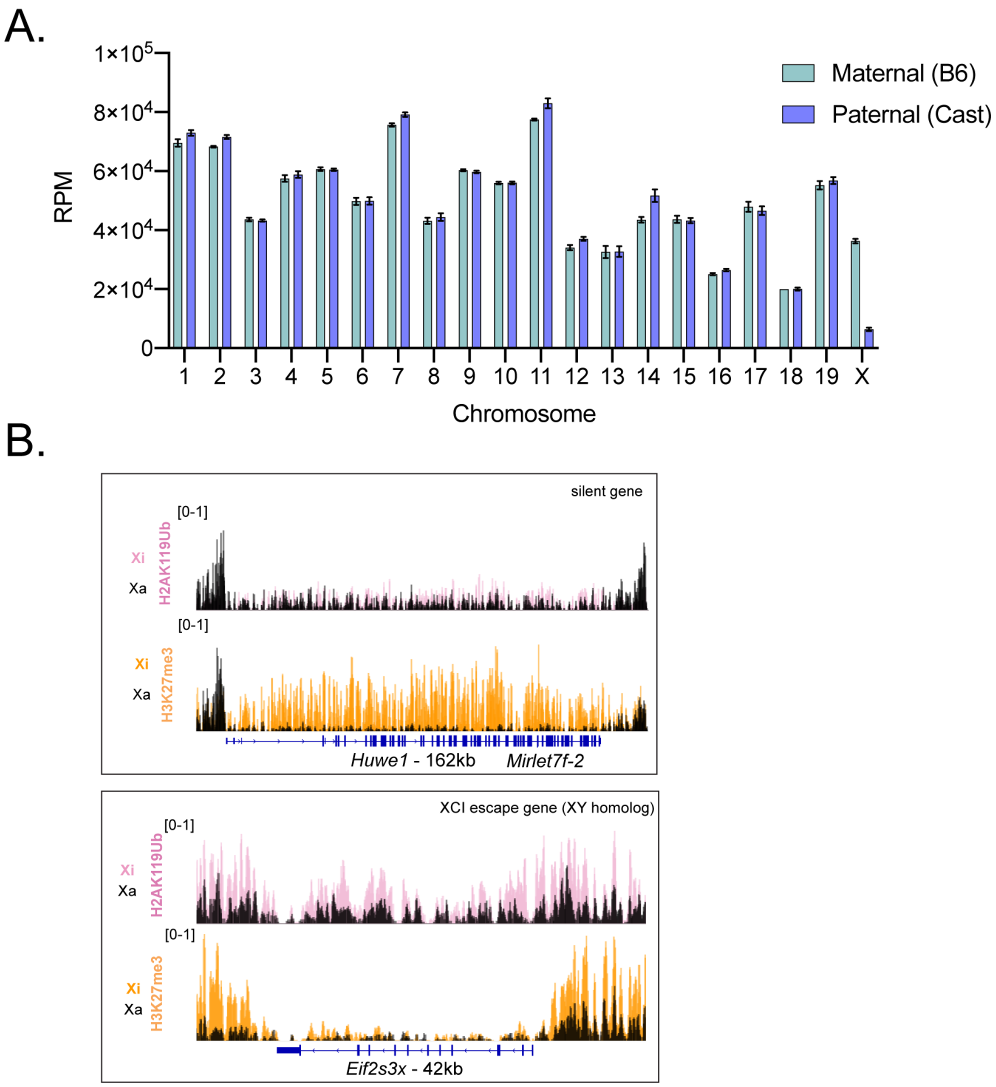


Supplementary Figure 1

**Supplementary Figure 1. A.** Reads per million (RPM) for all chromosomes in unstimulated T cells, separated by allele. The Y chromosome is not included as all samples were from female mice and there were no reads aligned to the Y chromosome. **B.** CPM-normalized tracks of H2AK119Ub (top) and H3K27me3 (bottom) enrichment at individual genes. Black represents enrichment on the Xa, while dark pink represents Xi enrichment of H2AK119Ub and dark orange represents Xi enrichment of H3K27me3. The genes shown are *Huwe1* (162kb), and *Eif2s3x* (42kb).

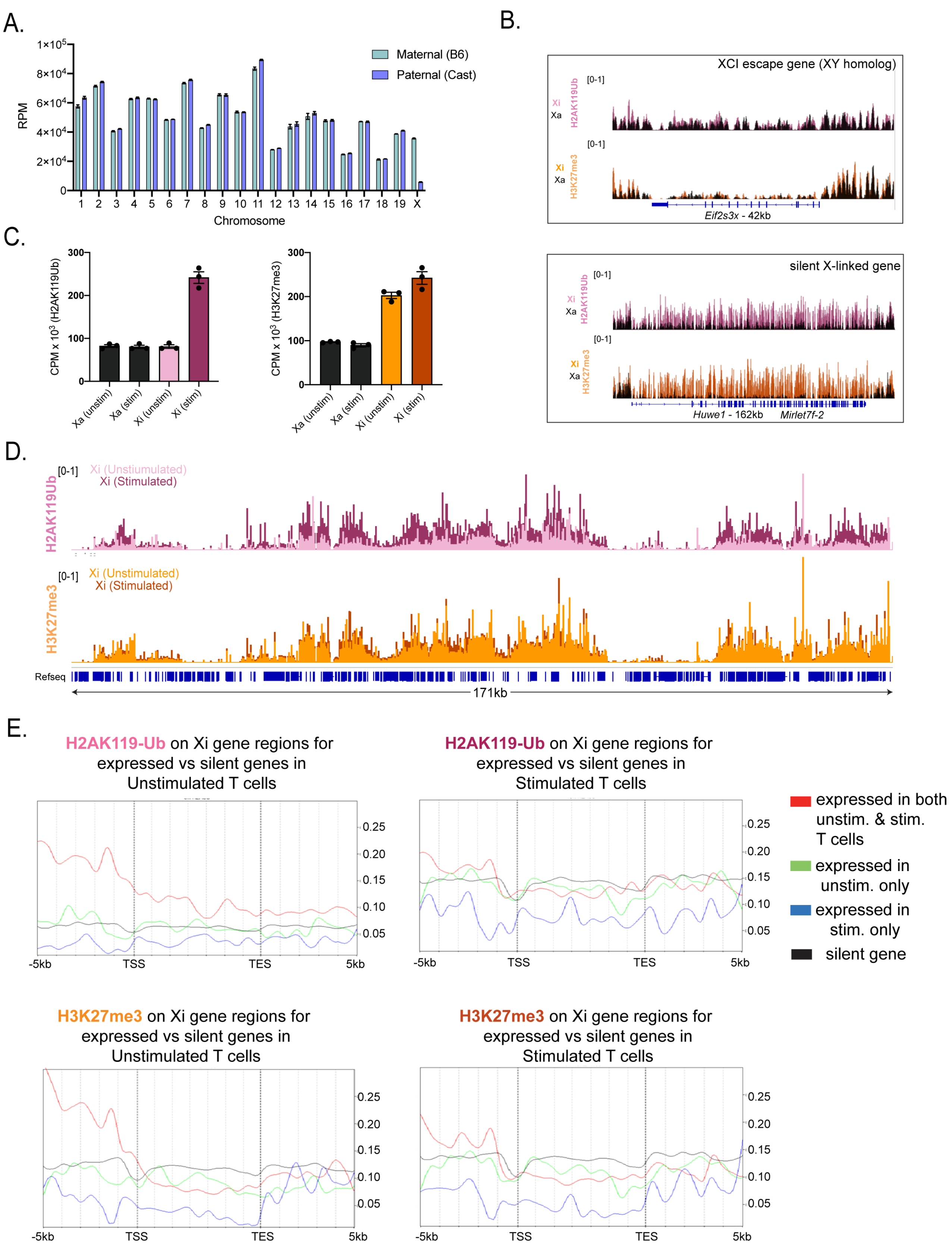


Supplementary Figure 2

**Supplementary Figure 2. A.** Reads per million (RPM) for all chromosomes in stimulated T cells, separated by allele. The Y chromosome is not included as all samples were from female mice and there were no reads aligned to the Y chromosome. **B.** CPM-normalized tracks of H2AK119Ub (top) and H3K27me3 (bottom) enrichment at individual genes. Black represents enrichment on the Xa, while dark pink represents Xi enrichment of H2AK119Ub and dark orange represents Xi enrichment of H3K27me3. The genes shown are *Huwe1* (162kb), and *Eif2s3x* (42kb). **C.** CPM of H2AK119Ub (left) or H3K27me3 (right) reads mapped to either the Xa or Xi in unstimulated or stimulated T cells. **D.** CPM-normalized track of X chromosome wide enrichment of H2AK119Ub (top) and H3K27me3 (bottom) in comparing Xi enrichment unstimulated (light pink and light orange) and stimulated (dark pink and dark orange) T cells. **E.** Metaplots show average enrichments +5kb upstream of the transcription start site (TSS) to -5kb downstream of the transcription end site (TES) separated by escape status (red- expressed in both unstimulated and stimulated T cells, green- expressed in unstimulated T cells, blue- expressed in stimulated T cells, black- subject to XCI.

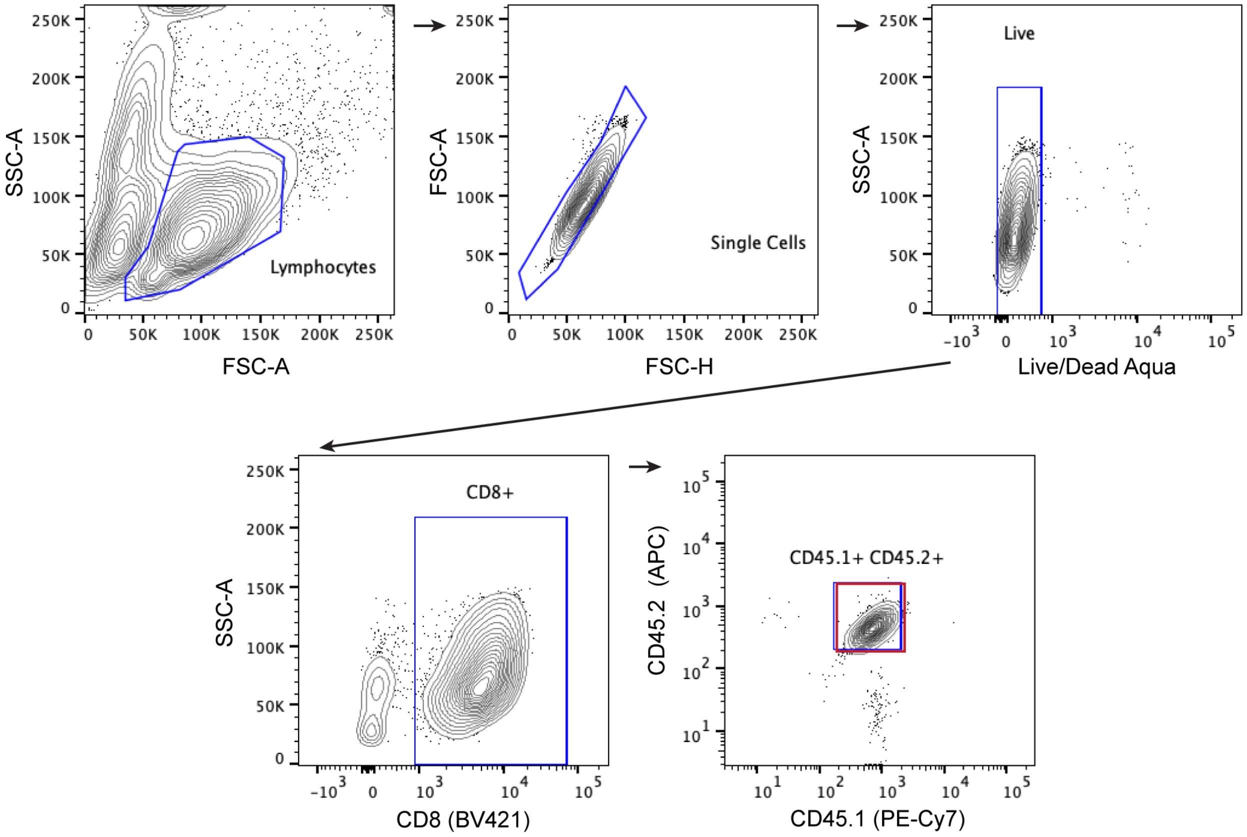


Supplementary Figure 3

**Supplementary Figure 3.** Gating strategy for OT-1 T cell sorts. Red box represents the sorted population.

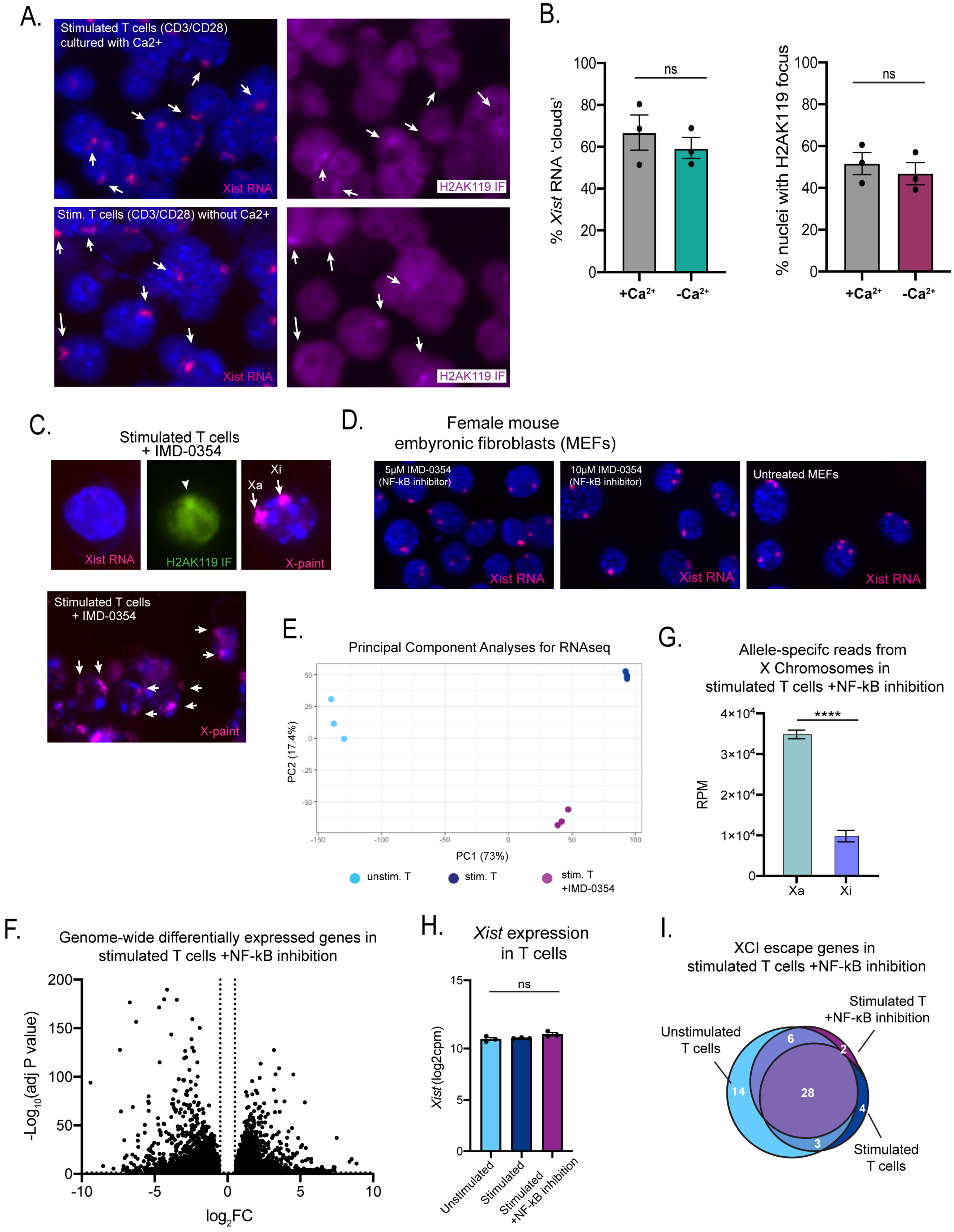


Supplementary Figure 4

**Supplementary Figure 4. A.** Representative fields (from 1 experiment) showing sequential *Xist* RNA FISH and H2AK119-Ub IF of $\alpha$CD3/$\alpha$CD28 stimulated T cells in the presence or absence of extracellular calcium for 72 hours. **B**. Quantification of *Xist* RNA clouds (left) and H2AK119-Ub foci (right) for $\alpha$CD3/$\alpha$CD28 stimulated T cells in the presence or absence of extracellular calcium for 72 hours. Averages of 3 independent experiments with SEM are shown with each dot representing a biological replicate. Statistical significance quantified using T test with Welch's correction. 87-118 nuclei (FISH) and 94-111 nuclei (IF) were counted per condition per experiment. **C.** Inset cells and representative field showing sequential *Xist* RNA FISH, H2AK119Ub IF, and X Chromosome DNA FISH (X-paint) for T cells stimulated with $\alpha$CD3/$\alpha$CD28 for 48 hours in the presence of 2.5μM IMD-0354. **D.** Representative fields of mouse embryonic fibroblasts (MEFs) treated with 5μM or 10μM of IMD-0354 for 2 days prior to *Xist* RNA FISH. **E.** Principal component analysis of RNA-seq samples from unstimulated T cells, $\alpha$CD3/$\alpha$CD28 stimulated T cells, and $\alpha$CD3/$\alpha$CD28 stimulated in the presence of 2.5μM IMD-0354 T cells. **F.** Volcano plot of genome-wide differentially expressed genes comparing T cells stimulated in the absence or presence of IMD-0354. **I.** Venn diagram of genes escaping XCI in unstimulated T cells, stimulated T cells, and stimulated in the presence of IMD-0354 T cells. **G.** Total reads per million (RPM) coming from the Xa and Xi alleles. Averages of 3 biological replicates with SD are shown. Statistical significance quantified using unpaired T test with Welch's correction, **** $p < 0.0001$. **H.** Log$_2$CPM values of *Xist* expression in T cells that were either unstimulated, stimulated, or stimulated in the presence of IMD-0354 from RNA-seq data set. Statistical significance quantified using a one-way ANOVA.

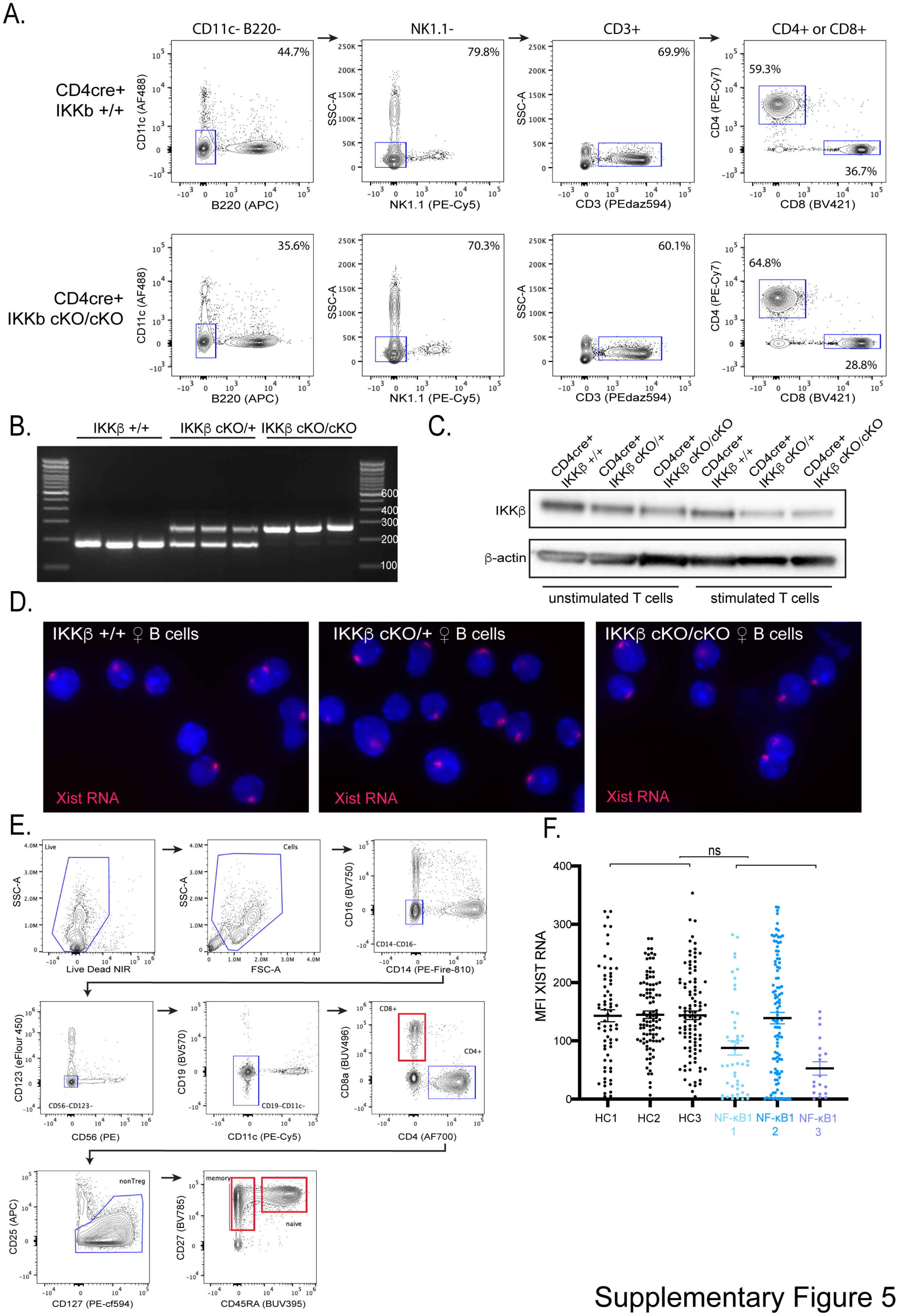


Supplementary Figure 5

**Supplementary Figure 5. A.** Representative flow plots of splenocytes isolated from cKO/cKO or WT littermate mice. Events were previously gated on singlets, live cells. **B.** Genotyping PCR gel for each of three IKKβ +/+, IKKβ cKO/+, and IKKβ cKO/cKO female mice, with expected amplification sizes of 200bp (floxed allele) and 180bp (WT allele). **C.** Western blot of unstimulated or αCD3/αCD28 stimulated $CD3^+$ T cells from IKKβ +/+, IKKβ cKO/+, and IKKβ cKO/cKO female mice. Blot was stained for IKKβ and β-actin. **D.** Representative fields (from 1 experiment) of *Xist* RNA FISH of CpG stimulated $CD23^+$ B cells from IKKβ +/+ (top), IKKβ cKO/+ (middle), and IKKβ cKO/cKO (bottom) female mice. **E.** Gating strategy for human PBMC cell sorts. Red boxes represents the sorted populations. **F.** Mean fluorescence intensity (MFI) of *XIST* RNA for sorted bulk $CD8^+$ T cells from healthy controls (black) and NF-κB1 patients (light blue, blue, purple) stimulated with αCD3/αCD28 for 48 hours, with each dot representing a nucleus. Statistical significance quantified using a Mann-Whitney non-parametric T test, with 69-103 nuclei (healthy) or 20-100 nuclei (patient) counted per sample. Due to low cell number, $CD8^+$ T cells from NF-κB1-4 were not isolated.

**Supplementary Table 1.** List of all genes escaping in unstimulated and stimulated T cells. Table includes escape status in unstimulated and stimulated T cells, as well as relevant expression information.

**Supplementary Table 2.** List of all expressed X-linked genes in unstimulated (tab 1) and stimulated T cells (tab 2). Table includes escape status in unstimulated and stimulated T cells, as well as biological replicate SRPM separated by allele.

**Supplementary Table 3.** List of all differentially expressed X-linked genes compared between stimulated T cells and T cells stimulated in the presence of 2.5μM IMD-0354. Table includes $Log_2$ Fold Change and adjusted P values, as well as escape status in = stimulated T cells and T cells stimulated in the presence of 2.5μM IMD-0354.

**Supplementary Table 4.** List of all *Xist* interactome genes differentially expressed in either unstimulated compared to stimulated T cells or stimulated T cells compared to T cells stimulated in the presence of 2.5μM IMD-0354. Table includes the average and standard deviation Z score for each stimulation condition, per differentially expressed gene.

**Supplementary Table 5.** Healthy and NF-κB1 patient information including age, sex, NFκB1 mutation (if applicable), clinical features (if applicable), and PBMC source information.
